# Chaos and repetition in algal evolution: A Chrysophyceae eco-genomic framework

**DOI:** 10.64898/2026.10.27.762293

**Authors:** Louis Graf, Emily Zajac, Ian Probert, Xiaoyu Zhang, Richard G. Dorrell

## Abstract

- The Chrysophyceae are an algal class that, within relatively short evolutionary distances, have repeatedly adopted multiple trophic strategies (photoautotrophy, mixotrophy, heterotrophy) and are ubiquitous across marine, freshwater and cryosphere ecosystems.
- Historical understanding of marine chrysophytes has lagged behind that of freshwater taxa, limiting our ability to identify what genes determine their cellular and ecosystem function.
- We leverage new transcriptomes, meta-genomes and computational predictions to reveal the functional, genetic and eco-physiological diversity of Chrysophyceae, including newly documented marine groups.
- We show that their common ancestor was likely a marine mixotroph, with multiple independent colonizations of freshwater and cryospheres, and evolution of photoautotrophic and heterotrophic life-strategies.
- We identify characteristic biogeographical distributions for different Chrysophyceae orders, including new ones linking marine and freshwater cryospheres.
- We document convergent gene losses relating to cryotolerance, photosynthesis, and phagocytosis, including novel families widely spread across the algal tree of life.
- We map meta-gene distributions of marker genes and impute a latitudinal shift in Chrysophyceae trophic strategy, with photoautotrophy predominant in polar oceans and heterotrophy in oligotrophic tropical waters.
- Our study identifies new gene markers that may be determinant of lifestyle shifts across the algal tree of life and reveals probable responses of marine chrysophytes to climate change.

## Introduction

Eukaryotic algae comprise more than 250,000 species (Frey, 2020; Kamiya, 2020; Guiry & Guiry, 2025). Algae are polyphyletic, with multiple evolutionary origins associated with the repeated primary and secondary endosymbiotic acquisitions of chloroplasts across the eukaryotes (e.g. Dorrell *et al*., 2017). They contribute substantially to global oxygen production (Friedlingstein *et al*., 2023) and forming the foundation of food webs (Sigee, 2005; Gasol & Kirchman, 2018).

Algal groups have radiated into diverse life strategies and cell forms. While most species are free-living and photosynthetic, others are symbiotic (Caron *et al*., 2009; Bjorbækmo *et al*., 2020; Chaffron *et al*., 2021). They have flexible feeding strategies, including photoautotrophy, obligate heterotrophy due to the loss of photosynthesis, and multiple forms of mixotrophy (e.g. Jeong *et al*., 2021a). This complicates algal roles in aquatic energy, carbon and nutrient cycles and in global biogeochemistry (Worden *et al*., 2004, 2015; Falkowski *et al*., 2008; Falkowski, 2012). Algal species may be unicellular, colonial, filamentous, free-living or sessile (Caron *et al*., 2012, 2017; Büdel *et al*., 2024), and vary dramatically in their fundamental biochemistry, e.g. in lipid metabolism (Guschina & Harwood, 2006; Lang *et al*., 2011; Li-Beisson *et al*., 2019). Finally, algae have adapted to virtually all illuminated habitats and ecological niches (e.g. De Vargas *et al*., 2015), including marine, freshwater, terrestrial and cryosphere ecosystems (Büdel *et al*., 2024), Nonetheless, many algal groups are confined to either marine or freshwater habitats, with only a few able to colonize the entire Earth ecosystem (Jamy *et al*., 2022).

The genetic mechanisms that determine algal ecological, physiological and metabolic traits remain poorly understood. Firstly, their trophic strategy and environmental adaptation are likely to be determined by multiple loci. Secondly, the deep evolutionary distances separating major algal clades erode the reliability of orthology-based functional inference. Finally, only a few algal species are experimentally tractable for reverse genetic characterization (e.g. *Chlamydomonas reinhardtii* [Chlorophyta], *Phaeodactylum tricornutum* [Heterokonts, Diatomae], *Nannochloropsis oceanica* [Heterokonts, Eustigmatophyceae], *Ectocarpus* sp.7 [Heterokonts, Phaeophyceae]). Consequently, algal genomes contain large numbers of genes with no characterized homologs (i.e. genomic “dark matter”), whose functions cannot be inferred by sequence similarity.

An alternative strategy to classify the functions of algal genes is to compare gene content across recently diverged groups in which specific traits evolved multiple times. The Chrysophyceae (within the Heterokontophyta) are ideal for this: despite radiating no more than 280-300 million years ago, their functional and ecological diversity mirrors that of algae at large (Brown & Sorhannus, 2010; Choi *et al*., 2024). The ancestor of Chrysophyceae possessed a plastid inherited from a secondary (or higher) red endosymbiotic event (Stiller *et al*., 2014; Dorrell *et al*., 2017), and was likely a single-celled facultative phago-mixotroph. However, extant Chrysophyceae include solitary, colonial, multicellular, free-living, and sessile forms, with some features (e.g., loricas) having evolved repeatedly (Andersen *et al*., 2025). Considering trophism, multiple ae genera (*Paraphysomonas; Spumella*; Grossmann *et al*., 2016; Jeong *et al*., 2021), have lost the ability to photosynthesize, returning to a state of obligate heterotrophy, while others (*Synura*) have lost phagocytosis and rely entirely on photosynthesis (Andersen, 2004; Kristiansen & Škaloud, 2017).

Previous studies of Chrysophyceae transcriptomes (notably Keeling *et al*., 2014; Beisser *et al*., 2017), have identified genetic trends associated with the loss of photosynthesis (Dorrell *et al*., 2019). These analyses have been based mainly on freshwater Chrysophyceae, with few described marine taxa (Andersen, 2004; Kristiansen & Škaloud, 2017). In parallel, metagenomic surveys have revealed that marine Chrysophyceae are diverse, abundant and ecologically relevant, including in poorly-studied polar oceans (Del Campo & Massana, 2011; De Vargas *et al*., 2015; Wideman *et al*., 2019; Delmont *et al*., 2022; Vaulot *et al*., 2022). Exploring the genetic features of marine Chrysophyceae is essential to understand their ecosystem functions, resolve their deep evolution, and to identify the genetic traits underlying their capacity to colonize all aquatic habitats.

Here, we exploit new transcriptomes and meta-genome assembled genomes from the global Earth ecosystem to identify genetic drivers of Chrysophyceae ecological and functional diversity. We show that marine Chrysophyceae are paraphyletic, with multiple highly diverse and uncultured clades that position their common ancestor as a marine phagomixotroph. We identify divergent biogeographical trends in different Chrysophyceae lineages, across cryosphere, temperate and tropical ocean biomes. We further identify new gene families correlated to phototrophic and phagotrophic lifestyles and cryosphere habitats that may be functional drivers of these traits. Finally, using quantitative metagenomics, we propose that marine Chrysophyceae transition from obligate photoautotrophy towards obligate heterotrophy with decreasing latitude, relevant to their future eco-physiology in heating oceans. Our study demonstrates how the microscale evolutionary diversity of algae can connect gene features, taxonomy, cellular behavior, and ecology at global scales.

## Material and Methods

### Sample collection, nucleic acid extractions and sequencing

We obtained ten cultured strains (*Glossomastix* sp. RCC3688, *Dinobryon* sp. RCC2294, *Paraphysomonas* sp. RCC5307, *Ochromonas* spp. RCC6673, RCC6677, RCC6719, RCC7845, RCC7846, RCC7855; *Ochromonas* sp. CCMP2296) representing poorly sampled marine Chrysophyceae and close relatives (Gérikas Ribeiro *et al*., 2020; Perrin & Dorrell, 2024). Cultures were maintained in K2 medium (Keller *et al*., 1987) made with ESAW (Berges *et al*., 2001) and repicked once a month, with RCC2294, RCC5307, RCC7845, RCC7846 and RCC7855 maintained at 4°C with constant light and RCC6673, RCC6677 and RCC6719 at 19°C with a 12h:12h light:dark cycle; both under cool white LEDs at 30-50 µE m^-2^ s^-1^ intensity.

RNA was extracted from 50mL of culture harvested by centrifugation (10min, 3450 rpm, 4°C) and washed with 1mL of PBS 1%. Phase separation was conducted with a 5:1 ratio of TRIzol™ (Invivogen, France) to chloroform (Sigma-Aldrich, Deutschland) and centrifugation (2min, 10.000rpm, 4°C). The supernatant aqueous phase was mixed with an equivalent volume of chloroform and centrifuged (2min, 10.000rpm, 4°C). Nucleic acids in the supernatant aqueous phase were precipitated with isopropanol (1h, -20°C), pelleted by centrifugation (30min, 10.000rpm, 4°C), washed with 100% ethanol, centrifugated (5min, 10.000rpm, 4°C), dried and resuspend in 50µL of RNAse-free H_2_O (Thermo Fisher, USA). Concentration and integrity of the nucleic acid was checked on a 1% agarose gel. When necessary, samples were treated with 1U of DNAse (Promega, France) per mg of RNA incubated (30min, 37°C) and RNA was precipitated with isopropanol (1h, -20°C), centrifugated (30min, 10.000rpm, 4°C), washed with 70% ethanol, centrifuged (5min, 10.000rpm, 4°C), dried and resuspended in 25µL of RNAse-free H_2_O. Concentration and integrity of the nucleic acid were checked using a Nanodrop photo-spectrometer and migration on a 1% agarose gel.

Sequencing was outsourced to Fasteris (CCMP2296, RCC3688, RCC6673, RCC6677) and Genewiz (RCC7845, RCC7846, RCC7855; Illumina MiSeq, 2×100bp), or BGI (RCC2294, RCC5307, RCC6719; DNBseq, 2×150bp). Raw data were deposited under NCBI SRA accession SAMN60274356-SAMN60274371.

### Transcriptome assembly and annotation

The following publicly available transcriptomes were processed along newly generated ones: *Chlorochromonas danica* SAG 933-7 (ERR13916157), *Chromulina chionophila* CCMP261 (SRR29150330), *Chrysamoeba radians* NIES-2890 (SRR27254668), *Chrysosaccus* sp. CCMP1156 (SRR29150319), *Hydrurus foetidus* (ERR2869477), *Kremastochrysopsis* sp. DR75b (SRR9878530), *Paraphysomonas bandaiensis* RCC383 (ERR12857270), *Paraphysomonas longispina (*SRR31641734, SRR31641735, SRR31641736)*, Pedospumella encystans* 1006 (SRR9203573) and *Poterioochromonas* sp. CCMP2060 (SRR29150312).

Reads were trimmed and filtered using fastp v0.24.1 (Chen, 2025) with default settings and the –detect_adapter_for_pe option, and *de-novo* assembled using rnaSPAdes v3.15.5 (Bushmanova *et al*., 2019) with default settings, and the main fast output (i.e. not hard or soft filtered) was used for subsequent analysis. For both newly assembled and publicly available transcriptomes, open reading frames (ORFs) were predicted using TIdeS v1.2.0 (Maurer-Alcalá & Kim, 2024) with the -p option, allowing partial ORFs, and the default database. Proteome completeness was assessed with BUSCO v 5.8.3 (Manni *et al*., 2021) with the “eukaryota_odb12” database, and translated ORFs were annotated using InterProScan v5.67-99 (Jones *et al*., 2014) with default options.

### Phylogenomic analysis

Newly annotated transcriptomes were combined with publicly available ones (MMETSP; Keeling *et al*., 2014), Metagenome Assembled Genomes (MAG) and Single-Cell Genomes (SCG) (Busi *et al*., 2022; Delmont *et al*., 2022), yielding 93 pteomes that represent the diversity of the Chrysophyceae, plus five outgroup taxa (Synchromophyceae and Pinguiophyceae; Supplementary Table 1). The 93 proteomes were processed with PhyloFisher v1.2.13 (Tice *et al*., 2021) following the workflow’s steps separately. Briefly, homologues to the 241 genes of the PhyloFisher database were identified in each library using hmmsearch v3.1 (Eddy, 2011). Candidate homologs were aligned along with multiple curation by PREQUAL (Whelan *et al*., 2018), MAFFT (Katoh *et al*., 2019), Divvier (Ali *et al*., 2019) and trimAl (Capella-Gutiérrez *et al*., 2009) with default settings. Single-gene trees were built using RAxML-NG 1.2.0 (Kozlov *et al*., 2019) under the LG4X+G4 model with 100 parametric bootstrap replicates, and inspected by eye using ParaSorter v.1.0.4 to correct orthologue and paralogue assignation. From the 241 orthologues, trees in which Chrysophyceae were polyphyletic were excluded, leaving 205, for which raw sequences were aligned as above and concatenated into a supermatrix containing 71,297 sites. Maximum likelihood reconstruction used IQ-tree v.3.0.0 (Wong *et al*., 2025), with an independent model for each partition selected automatically (-m MFP option) and 5,000 ultrafast bootstrap replicates (Hoang *et al*., 2018).

### Predictions of phago-mixotrophy using a gene-based model

First, closely-related taxa (predominantly MAGs) whose BUSCO coverage was deemed to be insufficient for functional trait prediction were merged using the PhyloFisher topology (>60% BUSCOs identified, <20% duplicated), following Dorrell *et al*. (2019). In total, 20 MAGs were combined into 9 merged MAGs (Supplementary Table 1), giving a final dataset of 82 proteomes (77 Chrysophyceae; 5 outgroups). These were tested for phagocytotic, photosynthetic, and prototrophic capabilities using the predictTrophicMode tool (Burns *et al*., 2018) run in default mode, identifying homologues to its 14,095 gene HMM models using hmmsearch v3.1 (Eddy, 2011), and predictions were visualized in R v4.5.1 (R Core Team, 2025). Manual corrections were made to three taxa. The non-pigmented *Cornospumella fuschlensis* AR4D6 retains most genes related to photosynthesis (Grossmann *et al*., 2016; Dorrell *et al*., 2019) and was predicted as phago-mixotroph, while *Pedospumella encystans* JBMS11 had low phagocytosis predictions (0.17 to 0.5) despite being shown experimentally to prey on bacteria (Boden *et al*., 2024), both were annotated as heterotrophic. Conversely, *Mallomonas* sp. CCAC0094 was predicted to be phago-mixotrophic by a small margin (0.41-0.6) despite no described phagocytosis in this genus (Andersen, 2004; Kristiansen & Škaloud, 2017), and was corrected to be photoautotrophic.

### Predictions of ancestral trait

Environmental, trophic and thermal traits were considered as discrete variables for the reconstruction of ancestral states. Habitat was coded as saline, freshwater or soil, according to the environment where each strain was collected, or eDNA assembled into MAGs was sampled (Supplementary Table 1). Trophism was coded in four states according to proteome-based predictions (Burns *et al*., 2018): photoautotrophs, capable of photosynthesis only; mixotrophs, capable of photosynthesis and phagocytosis and/or prototrophy; heterotrophs, capable of phagocytosis and/or prototrophy only; and parasites, lacking any predicted autonomous trophic ability (Supplementary Figure 1). As measured thermal tolerance was not directly assessed for most taxa, we coded the thermal traits according to collection site: polar habitats were classified as cryosphere-associated and others as mesosphere-associated. *Tara* MAGs were considered marine, with the ocean basin of greatest mapped read depth used as an indicator of biogeography, e.g. those marked “ARC” (Arctic) assigned to the cryosphere (Delmont *et al*., 2022). Glacier meltwater *Hydrurus* MAGs assembled by Busi *et al*., (2022) were annotated as cryosphere and freshwater.

Ancestral states were reconstructed using the “ace” function from the R package “ape” (Paradis & Schliep, 2019) with maximum likelihood models, comparing equal rates (“ER”), all-rates different (“ARD”) or symmetrical (“SYM”) models using the Akaike Information Criterion. The symmetrical model performed better for environmental traits and the equal rates model for the trophic and thermal traits. Reconstructions were plotted as pie charts on the nodes of the phylogenomic tree using iTOL (Letunic & Bork, 2024).

### Biogeography

rRNA sequences were identified in newly generated and re-assembled transcriptomes using barrnap (https://github.com/tseemann/barrnap). Due to their assembly methodology; MAGs do not contain rRNA sequences (Delmont *et al*., 2022) but we were able to identify rRNA in the three SCGs. These sequences were combined with publicly available sequences (ncbi consulted on October 21^st^ 2025) representing the cultivable diversity (i.e. excluding “uncultured” or “environmental” sequences) of the Chrysophyceae for a total of 226 Chrysophyceae and 9 outgroup sequences. Sequences were aligned using MAFFT v7.526 (Katoh *et al*., 2019) using --auto settings and adjusted by eye in AliView v1.28 (Larsson, 2014). The alignment was used for phylogenetic reconstruction using IQ-tree v.3.1.1 (Wong *et al*., 2025) with the GTR+F+R6 model (selected by -m MFP) and 1,000 ultrafast bootstrap replicates (Hoang *et al*., 2018). The reconstruction was constrained using the topology of the multigene phylogenomic topology (-g option; Supplementary Figure 1).

All V4 barcode ASVs assigned to Chrysophyceae and Synurophyceae (1,259 in total) were recovered from the Ocean Barcode Atlas webserver (Vernette *et al*., 2021; https://oba.mio.osupytheas.fr/ocean-atlas/OBA_analyse) accessed on October 22^nd^ 2025, using default (clade taxonomy) parameters. The ASVs were aligned with the 235 reference rRNA sequences using MAFFT v7.526 (Katoh *et al*., 2019), adjusted by eye in AliView v1.28 (Larsson, 2014) and trimmed to the V4 region. The alignment was used for phylogenetic reconstruction using IQ-tree v.3.1.1 (Wong *et al*., 2025) with the GTR+F+I+R8 model (selected by -m MFP) and 1,000 ultrafast bootstrap replicates (Hoang *et al*., 2018). The reconstruction was constrained using the topology of the reference rRNA topology (-g option; Supplementary Figures 2).

The tree topology was used to determine the taxonomic affiliation of the ASVs according to their relationship to known cultivable sequences (Supplementary Figure 2; Supplementary Table 2). Because different *Tara* missions and stations used different filter sizes (Pesant *et al*., 2015), these were grouped into size fractions: “pico” (“0.22-1.6µm”, “0.22-3µm”, “0.8-3µm”, “0.8-5µm”), “nano” (“0.8-20µm”, “5-20µm”), “micro” (“20-180µm”), “macro” (“180-2000µm”) and “unfiltered” (“>0.8µm”, “>3µm”). Mean abundances for each size fraction and depth (surface, SRF; Deep Chlorophyll Maximum, DCM) were plotted on world maps using the Mollweide projection. Principal Component Analyses (PCA) were performed with *Tara* physico-chemical variables of each stations (Pesant *et al*., 2015), with missing values imputed using imputePCA from the missMDA package (Josse & Husson, 2016). The mean ASV abundances were plotted on the PCA as supplementary variable. All calculations and plotting were conducted in R v4.5.1 (R Core Team, 2025).

Finally, habitat models for each marine Chrysophyceae group were constructed from *Tara* V4 surface abundances (Supplementary Figure 2) following Penot-Raquin et al. (2026), using a Random Forests approach with the R package tidymodels (https://www.tidymodels.org) with an 80/20 training/testing ratio, and hyperparameters for the numbers of trees and min_n estimated and adjusted for each group (Supplementary Table 3). Environmental parameters were downloaded from the World Ocean Atlas 2018 (Boyer *et al*., 2018) at 1° resolution, using objectively analyzed means over 2005-2017 when available.

### Comparative genomics

Genomic diversity of 68 single or merged Chrysophyceae with >40% BUSCO coverage was explored over four dimensions for binary categories (presence/ absence) represented by more than 15 single or merged. These were: two predicted lifestyle traits (photosynthesis, phagocytosis), and two sampled habitats (salinity, cryosphere). For photosynthesis and phagocytosis, proteomes were classified as photosynthetic vs. heterotrophic and phagotrophic vs. photoautotrophic; respectively; according to the trophic lifestyles predicted by genome-scale modelling (Supplementary Table 1). For trophism analyses, taxa with identified discrepancies between genome)-scale predictions and observed lifestyles (*Cornospumella fuschlensis* AR4D6, *Pedospumella encystans* JBMS11, *Mallomonas* sp. CCAC0094, see above), leaving 65 proteomes. For environmental and thermal trait analysis, proteomes were classified as freshwater or marine and cryosphere or mesosphere according to collection environment as above, with soil taxa assigned to freshwater (Supplementary Table 1).

PFAM annotations were compared using a two-sided Fisher test based on the presence/absence of each PFAM recovered from the InterProScan v5.67-99 annotation (see above), and a two-sided Wilcoxon test on the proportion of each PFAM in the total proteome of each taxon, normalized by BUSCO completeness. For both tests, *p-*values were adjusted in R using the false discovery rate method (Benjamini & Hochberg, 1995), and PFAMs were considered significant if both tests reached an adjusted p-value < 0.05.

### Biogeographical analysis of gene families

To explore the function of oceanic Chrysophyceae, the relative abundance and expression of three representative gene families were mapped: PF01716 (*psbO*), PF11999 (IBPs), and PF14845 (beta-acetyl hexosaminidase). PF01716 and PF11999 were selected due to their prior use as meta-genomic markers for photosynthesis (Pierella Karlusich *et al*., 2023) and cryophily (Dorrell *et al*. 2023) respectively, with 168 and 36 MATOUs phylogenetically reconciled to Chrysophyceae PF01716 and PF11999, respectively, extracted from published trees.

PF14845 was selected due to its strong enrichment in phagocytotic taxa (i.e., absence from obligate photoautotrophs; see above). PF14845-containing sequences were retrieved from all 77 Chrysophyceae proteomes (130 total sequences) and 79 proteomes covering the diversity of the ToL (190 total sequences; Novák Vanclová *et al*., 2024). Homologues were, aligned using MAFFT v7.526 (Katoh *et al*., 2019) using --auto and manually trimmed to remove highly divergent regions. A phylogenetic tree was constructed using FastTree v2.2.0 (Price *et al*., 2009) using the -lg -cat 20 -nome options and its topology was transformed to a binary constraint with the TreeToConstraint.pl script (https://microbesonline.org/fasttree/constrained.html).

An HMM built from the aligned cultured species sequences using hmmerbuild v3.1 (Eddy, 2011), and was searched against all sequences in the MATOU-v1.5 dataset (Pierella Karlusich *et al*., 2025) using hmmsearch v3.1 with a domain e-value cut off of 10^-20^. Candidate MATOUs were filtered by a reciprocal MMseqs2 easy-search (Steinegger & Söding, 2017) with e-value threshold 10^-30^ against the 79 reference proteomes, excluding those whose best hits were not related to PF14845-containing sequences, leaving 6,885 MATOUs. These were aligned to the 320 Chrysophyceae and reference sequences using MAFFT v7.526 with --auto, and a tree was constructed with FastTree v2.2.0 with the -lg -cat 20 -nome options and constrained with the topology of the Chrysophyceae and reference PF14845 tree topology.

Finally, 161 MATOUs that reconciled as monophyletic with Chrysophyceae sequences (resolved in 4 polyphyletic clades) were extracted from the constrained tree. Their relative abundances in *Tara* meta-transcriptome (metaT) and meta-genome (metaG) data were normalized to the total relative abundance of all sequences annotated as Chrysophyceae in the DADA2 classification of the MATOU-v1.5 dataset. Normalized metaT and metaG abundances were mapped, and PCA and distribution predictions made following the ASVs methods above.

## Results

### Phylogenomic tree of the Chrysophyceae

We assembled a collection of proteomes assembled from *de novo* sequenced and publicly available transcriptomes and annotated MAGs (Keeling *et al*., 2014; Beisser *et al*., 2017; Grigoriev *et al*., 2021; Busi *et al*., 2022; Delmont *et al*., 2022). These covered the ecological and functional diversity of the Chrysophyceae, including 34 freshwater and 50 marine taxa, 18 taxa from cryosphere sites, and multiple trophic lifestyles (Supplementary Table 1). To provide an evolutionary framework to study the diversification of the Chrysophyceae we used an alignment of 205 orthologs (71,297 amino acid positions) to reconstruct a robust phylogenetic tree (Figure 1a).

**Figure 1.**
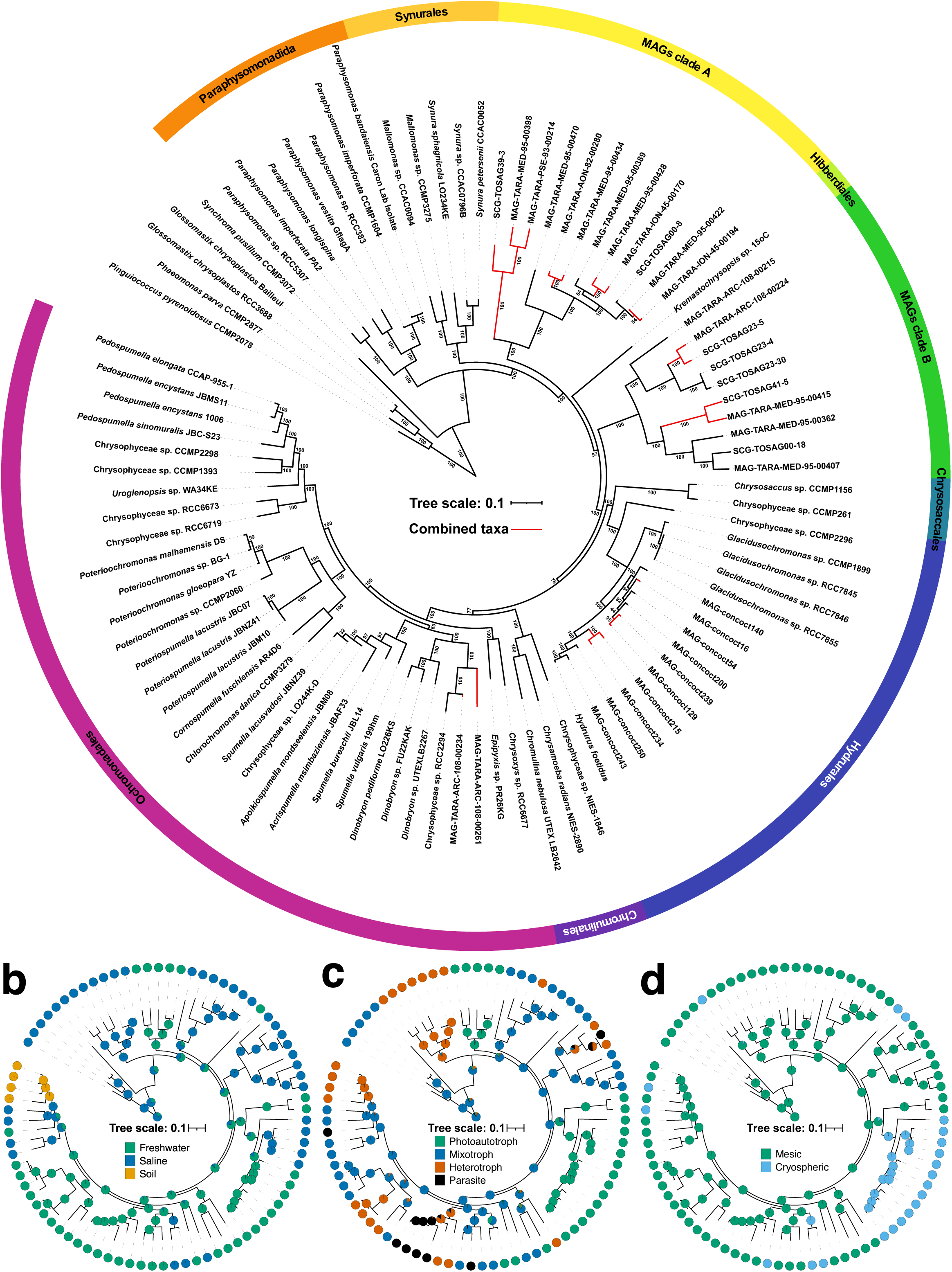
Evolutionary framework of the Chrysophyceae. **a.** Maximum-likelihood phylogenetic reconstruction of the class Chrysophyceae based on the concatenation of 205 orthologues. The tree was inferred with IQ-Tree v 3.1.1 using independent models for all partitions. Statistical supporting values were assessed by Ultrafast Bootstrap Approximation and are shown near the nodes. **b-d.** Ancestral traits reconstruction. Pie charts on nodes indicate the most likely state of the trait of that ancestor, as predicted by maximum likelihood models (see methods). **b.** Reconstruction of environmental traits coded as saline, freshwater or soil. **c.** Reconstruction of trophic traits coded as photoautotrophs, mixotrophs, heterotrophs, or parasites. **d.** Reconstruction of thermal traits coded as cryosphere-associated or mesosphere-associated.

Our tree was consistent with the known Chrysophyceae taxonomy (e.g. Dorrell *et al*., 2019; Bock *et al*., 2022). The Paraphysomonadida lineage diverged first (Ultrafast Bootstrap [UFB] 100%), followed by the Synurales (UFB 100%) and monophyletic Hibberdiales, Hydrurales, Chromulinales and Ochromonadales, all with full support (UFB 100%). All described genera were recovered as monophyletic (Figure 1a), except *Spumella*, which was paraphyletic to the genera *Acrispumella* and *Apoikiospumella* (UFB 100%).

Outside of the cultured taxa, the Metagenomes Assembled Genomes (MAGs) and Single-Cell Genomes (SCGs) formed four distinct and highly-supported lineages (Figure 1a; UFB 100%). Two of these (i.e. MAG clades A and B) were composed exclusively of uncultured marine taxa and branched after the Paraphysomonadida and Synurales, separated by a single freshwater snow alga, *Kremastochrysopsis* sp. DR75b (Figure 1a; UFB 97%; Raymond & Remias, 2019). The other two uncultured lineages were paraphyletic, including previously described taxa related to cryosphere habitats. Within the order Hydrurales, *Hydrurus foetidus* was nested among ten MAGs (clade C) assembled from glacier-fed streams (Busi *et al*., 2022), sister to marine strains isolated from the Arctic and Antarctic oceans (CCMP2296, CCMP1899, RCC7845, RCC7846, RCC7855; Figure 1a). A second lineage was composed of two marine MAGs (clade D) grouped with an Arctic strain (RCC2294), provisionally named *Dinobryon faculiferum* (Figure 1a; UFB 100%). We note that this annotation based on phylogeny does not reflect the understanding of *Dinobryon* morphology and ecology (as a colonial and freshwater genus), and this clade formed a sister to the freshwater strains *D. pediformis*, UTEX-LB-2267 and FU22KAK.

### Evolution of Chrysophyceae functional traits

Using our phylogenomic tree, we reconstructed the ancestral states of important traits that vary across Chrysophyceae taxa (Figure 1b-d). We focused on three traits: environmental (i.e. freshwater, soil or saline taxa), trophic (i.e. photoautotroph, heterotroph or phagomixotroph) and thermal (i.e. cryosphere or mesosphere).

Mapping of the environmental traits revealed that only the soil Chrysophyceae (*Pedospumella*) formed a monophyletic environmental group (Figure 1b). Reflecting the distribution of known freshwater taxa (*Poterioochromonas, Poteriospumella*, *Dinobryon*, Synurales), most ancestral states computed for late-diverging Chrysophyceae families were freshwater (i.e. Ochromonadales 94 %; Chromulinales 97.8%; Hydrurales 79.4%, and the common ancestor of all these groups 87.2%). Similarly, reflecting that *Paraphysomonas* is a marine genus, the common ancestor of Paraphysomonadida was predicted to be saline (87.2%). We nonetheless identified multiple marine and freshwater taxa were sister to one another, indicating repeated transitions across the salinity barrier (Figure 1b). Factoring in the newly discovered marine diversity and deep branching positions of the MAG clades A and B, a saline origin was predicted marginally for the Chrysophyceae last common ancestor (53.3%, Figure 1b).

Next, we analyzed the evolution of predicted trophic lifestyles (Figure 1c). To analyse predict trophic modes of uncultured taxa, we applied a comparative genomic predictive model (Burns *et al*., 2018) across the complete dataset, using cultured taxa with known lifestyles as controls (Supplementary Table 1; Supplementary Figure 3-4). Three taxa were manually corrected due to discrepancy between model predictions and cellular observation (Materials & Methods, Supplementary Table 1; Supplementary Figure 3-4). A further 11 taxa predicted to be parasitic or osmotrophs (i.e. lacking both photosynthesis and phagocytosis) all had very low BUSCO coverage (4.7%-40.8%; Supplementary Table 1), suggesting a prediction resulting from incomplete assemblies rather than biology. Of the 65 taxa with robust assignments (excluding “parasites” and “osmotrophs”), 47 were phagotrophic and 18 obligate photoautotrophs, while 45 were photosynthetic and 20 obligate heterotrophs.

None of the marine MAGs was predicted to be an obligate photoautotroph (Supplementary Figure 3-4). Both MAG clades A and B contained predicted phago-mixotrophs and obligate heterotrophs lacking photosynthesis (Figure 1c; Supplementary Figure 3-4), whereas MAG clade D and freshwater MAGs were predominantly obligate photo-autotrophs. Ancestral reconstructions suggested that the common ancestor of the Chrysophyceae was likely a phago-mixotroph (80.1%), with obligate photoautotrophs and heterotrophs originating via secondary losses of these traits. Five ancient transitions to obligate photoautotrophy were identified in the ancestors of Synurales, Chrysosaccales-Hydrurales, *Chrysamoeba, Epipyxis* and MAG clade C, and four to heterotrophy in the common ancestors of *Paraphysomonas, Pedospumella, Poteriospumella* and *Spumella* and within MAG clade B (Figure 1c).

Finally, the common ancestor of the Chrysophyceae was predicted to be mesophilic mesosphere (i.e., from a non-glacier and non-polar habitat) with high probability (99.9%). The 19 cryosphere taxa were distributed in seven monophyletic groups, most of them (13, including both freshwater snow and polar oceans) within the Hydrurales (Figure 1d).

### Biogeography of marine Chrysophyceae

Next, we explored the spatial distributions of different marine Chrysophyceae groups, taking advantages of the large, automatically comparable metabarcoding datasets and extensive, standardized environmental contextual data of the *Tara* Oceans expeditions. Chrysophyceae 18S rRNA V4 ASVs from the *Tara* Ocean Barcode Atlas (Vernette *et al*., 2021) were mapped for individual depth and size fraction combinations (Supplementary Figure 5a), related to physical and pigment variables by PCA (Supplementary Figure 5b-c), and projected globally, including oceanic regions not covered by *Tara*, via a Random Forests approach (Supplementary Figure 5d).

Chrysophyceae were detected in all oceans and at all size fractions. The greatest relative abundances were identified in the “Pico” fraction of <5 µm (mean ± standard deviation: 0.01135 ± 0.02124), approximately five times more than in the “Nano” fraction of <20 µm (0.00245 ± 0.00586); and 15 times more than in the “Micro” fraction of <200 µm (0.00075 ± 0.00371) and the “Macro” fraction of <2000 µm (0.00085 ± 0.00725; pairwise Wilcoxon *p-*values < 10^-05^ Supplementary Figure 6). Surface abundances (0.00696 ± 0.0148) were generally higher than at the DCM (0.00425 ± 0.0147) but not significantly (Wilcoxon *p*-value 0.4653). Furthermore, we noted that “Nano” and “Micro” size fractions were relatively more abundant in polar stations, and a large “Pico” size fraction was recorded at one DCM station in the equatorial Pacific (station 041; Supplementary Figure 5a). Class-level abundance was weakly, but not significantly, positively associated with temperature, salinity, pH, iron and CO_3,_ particularly in surface depths (Supplementary Figure 5b-c). Accordingly, random forest projections suggested Chrysophyceae are ubiquitous across the ocean, albeit with lower relative abundance in the Northern Atlantic than other regions (Supplementary Figure 5d).

### Distributions of individual Chrysophyceae groups

Next, to investigate if different taxa have different biogeographical distributions, we classified the ASVs using a constrained phylogeny based on annotated Chrysophyceae 18S rDNA sequences (Supplementary Figure 2). We identified 23 monophyletic lineages: 15 related to known cultivable diversity, five related to MAGs, and three (hereafter “environmental clades I, II and III”) branching within the Hibberdiales and Chrysosaccales but without cultivable or MAG representatives (Supplementary Figure 2).

The largest proportion of total Chrysophyceae relative abundance (summing all stations, depths and size fractions) was recovered in MAG clades A and B (24.1% and 27.01%, respectively), followed by a clade sister to the genus *Paraphysomonas* (16.45%), potentially corresponding to *Paraphysomonas formanifera* and the freshwater isolates JBC29 and JBNZ40 (Boenigk et al., 2005), and the *Dinobryon*-like clade (12.34%) including “*Dinobryon” balticum* and the polar strains RCC2290 and RCC2294. Cultivable taxa were less abundant (*Paraphysomonas* [7.51%]; *Pedospumella* [3.82%]; *Poterioochromonas* [2.23%]). Conversely, we noted genera not detected in *Tara* ASVs (e.g. *Spumella*, *Uroglena*) and confirmed the freshwater nature of the order Synurales (with only one single ASV, 175925, in a single station).

Distinct biogeographical patterns were observed for the nine higher taxonomic Chrysophyceae ranks (Figure 2; Supplementary Figure 7-24). Despite different total relative abundances, the orders Segregatales, Hibberdiales, and Ochromonadales shared broadly similar distributions with elevated abundances in polar and subpolar regions and lower ones toward the tropics (Figure 2a-c). The Segregatales showed the sharpest predicted bipolar signal, with continuous high abundance across the Southern Ocean and the Arctic/ North Atlantic and near-zero abundance throughout subtropical and tropical gyres (Figure 2a). Despite this distribution, the abundance of the Segregatales was not correlated to temperature, salinity or latitude, correlating positively to pigments (chlorophyll c3, chlorophyll a, peridinin, hexanoylfucoxanthin, Supplementary Figure 16). The bipolar signal of Hibberdiales was less marked, with moderate abundances projected outside the poles and no consistent environmental correlation (Figure 2b; Supplementary Figure 19). Ochromonadales were the most broadly distributed and abundant, combining polar/subpolar peaks with relatively elevated abundance across low-to-mid latitudes and local maxima in coastal regions (Figure 2c). They were positively correlated to inorganic nutrients (NO_3_, HCO_3_, O_2_) at the surface but negatively at the DCM (Supplementary Figure 24). Ochromonadales sublineages differed, with the *Dinobryon-*like lineage bipolar, *Pedospumella* tropical, and *Poterioochromonas* ubiquitous but most abundant in Indian and Pacific open oceans (Supplementary Figure 25a).

**Figure 2.**
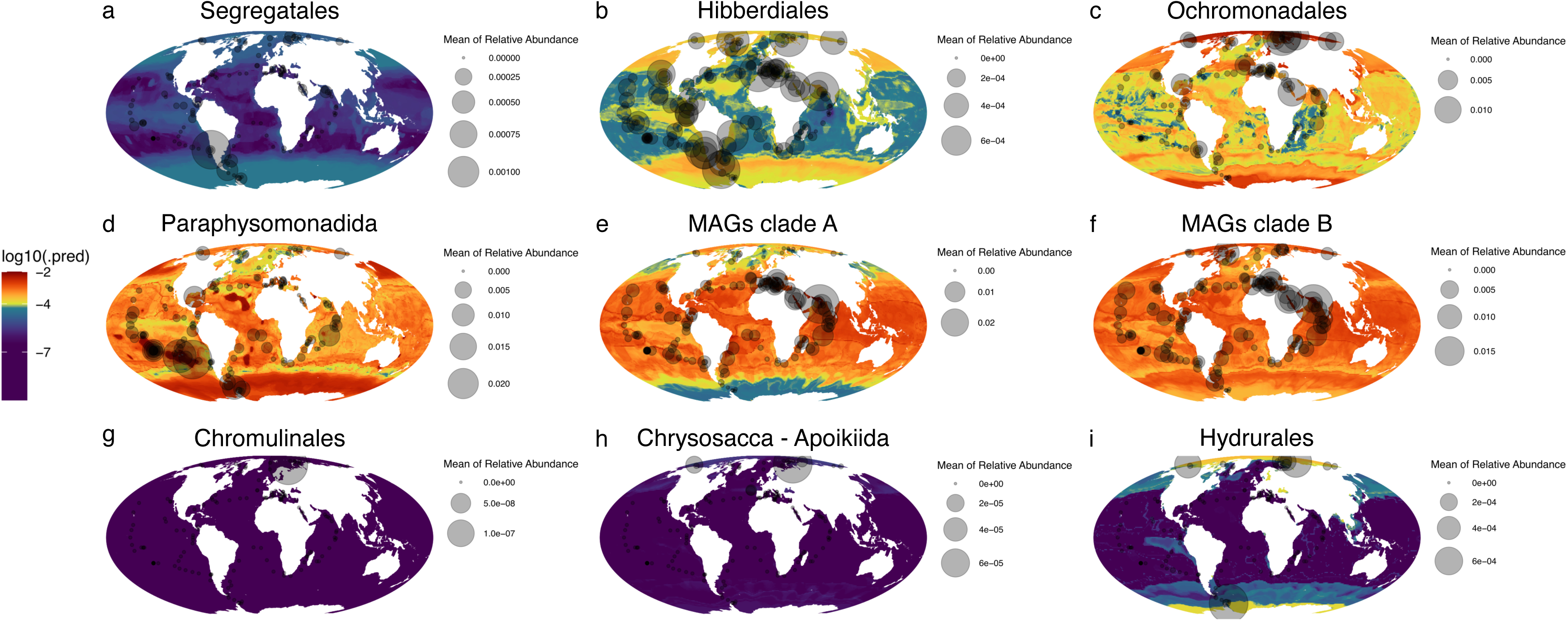
Predicted abundances at the surface of the Chrysophyceae orders at the global scale. Metabarcodes of the Chrysophyceae were phylogenetically assigned to the different orders (Supplementary Figure 2; Supplementary Table 2) and their relative abundances at the surface were used to predict distributions using a random tree forest approach and satellite data. **a.** Segregatales. **b.** Hibberdiales. **c.** Ochromonadales. **d.** Paraphysomonadida. **e.** MAGs clade A. **f.** MAGs clade B. **g.** Chromulinales. **h.** Chrysosaccales. **i.** Hydrurales.

Paraphysomonadida, including its highly abundant sister-clade, were widely distributed, in particular in tropical, subtropical, and temperate oceans (Figure 2d), with the greatest projected abundances in subregions (e.g., the Persian Gulf) with temperatures > 30°C (Pous *et al*., 2015). High relative abundances were also observed and projected in subpolar regions of the Southern Ocean, North Atlantic and Pacific, although Paraphysomonadida abundances never dropped under 10^-04^. The different Paraphysomonadida sublineages showed various distributions, with *Paraphysomonas* itself most abundant in subpolar regions, its highly abundant sister-lineage ubiquitous but particularly frequent in the Subtropical Pacific and Atlantic Oceans, and *Lepidochromonas* largely restricted to tropical/equatorial regions (Supplementary Figure 25b). PCAs detected no strong environmental correlation for Paraphysomonadida or its sublineages (Supplementary Figure 17).

MAG clade A was predominantly associated with subtropical and tropical oceans, particularly central and phosphate-limited regions (Figure 2e), with the highest abundances in Mediterranean and Red Seas stations (e.g. stations 023, 048). Much lower abundances were predicted and observed in temperate and subpolar latitudes, upwelling zones and coastal regions (Supplementary Figure 18). At both depths, abundances were positively correlated to temperature and salinity and negatively to inorganic nutrients (NO_3_, PO_4_).

MAG clade B was abundant and ubiquitous (Figure 2f), including in tropical Red Sea and Indian Ocean stations but also in subpolar regions and lacked clear latitudinal transitions. Nonetheless PCAs showed weak positive correlations to temperature and salinity at the surface and weak negative correlations to NO_3_ at both depths (Supplementary Figure 20).

Hydrurales were strikingly and almost exclusively concentrated in polar and subpolar regions, with the highest abundances around Antarctica and in the Arctic Ocean (Figure 2i; Supplementary Figure 21). The transition from high to low abundance was extremely sharp, occurring over a relatively narrow latitudinal band, and Hydrurales were virtually absent across the tropical and subtropical ocean and in stations with temperatures > 6°C. Accordingly, surface and DCM abundances showed clear negative correlations to temperature and salinity, and positive ones to signed latitude and inorganic nutrients (Supplementary Figure 24).

Finally, the orders Chromulinales and the *Chrysosacca-Apoikiida* clade showed globally very low and patchy abundances that preclude robust biogeographic interpretation (Figure 2g-h).

### Comparative genomics

Next, we linked this diversity to gene content using PFAM annotations, a portable and widely-used resource for identifying lifestyle- and environment-specific gene families in eukaryotic algae (Mistry *et al*., 2021; Dorrell *et al*., 2023). We reported PFAMs that tested with corrected *p*-value < 0.05 in tests both of presence/ absence and relative occurrence (see Materials and Methods).

#### Photosynthetic versus heterotrophic taxa

233 PFAMs were significantly enriched in either photosynthetic or heterotrophic taxa (Figure 3a; Supplementary Table 3). Most (188) were exclusive to or enriched in photosynthetic taxa, suggesting net gene loss in obligately heterotrophic species (Supplementary Table 3). These included 25 relating to photosynthesis and photosystem biogenesis, including photosystem subunits, assembly factors, chlorophyll a/b-binding proteins, chloroplast protein complexes, chloroplast translation activators, and Rubisco assembly chaperones. A further 12 related to pigment biosynthesis, (chlorophyll via the tetrapyrrole pathway, and carotenoids via the plastidial mevalonate pathway), 12 to metal and cofactor homeostasis, and 7 to plastidial amino acid synthesis (lysine, aromatic and branched-chain amino acids; Dorrell *et al*., 2019), alongside a mitochondrial ornithine carbomoyltransferase (PF02729) implicated in diatoms to be involved in plastidial amino acid recycling (Levering *et al*., 2016; Nonoyama *et al*., 2019). 45 PFAM were enriched in heterotrophic taxa (Supplementary Table 3), including 7 involved in protein degradation, which could relate to phagocytotic activity.

**Figure 3.**
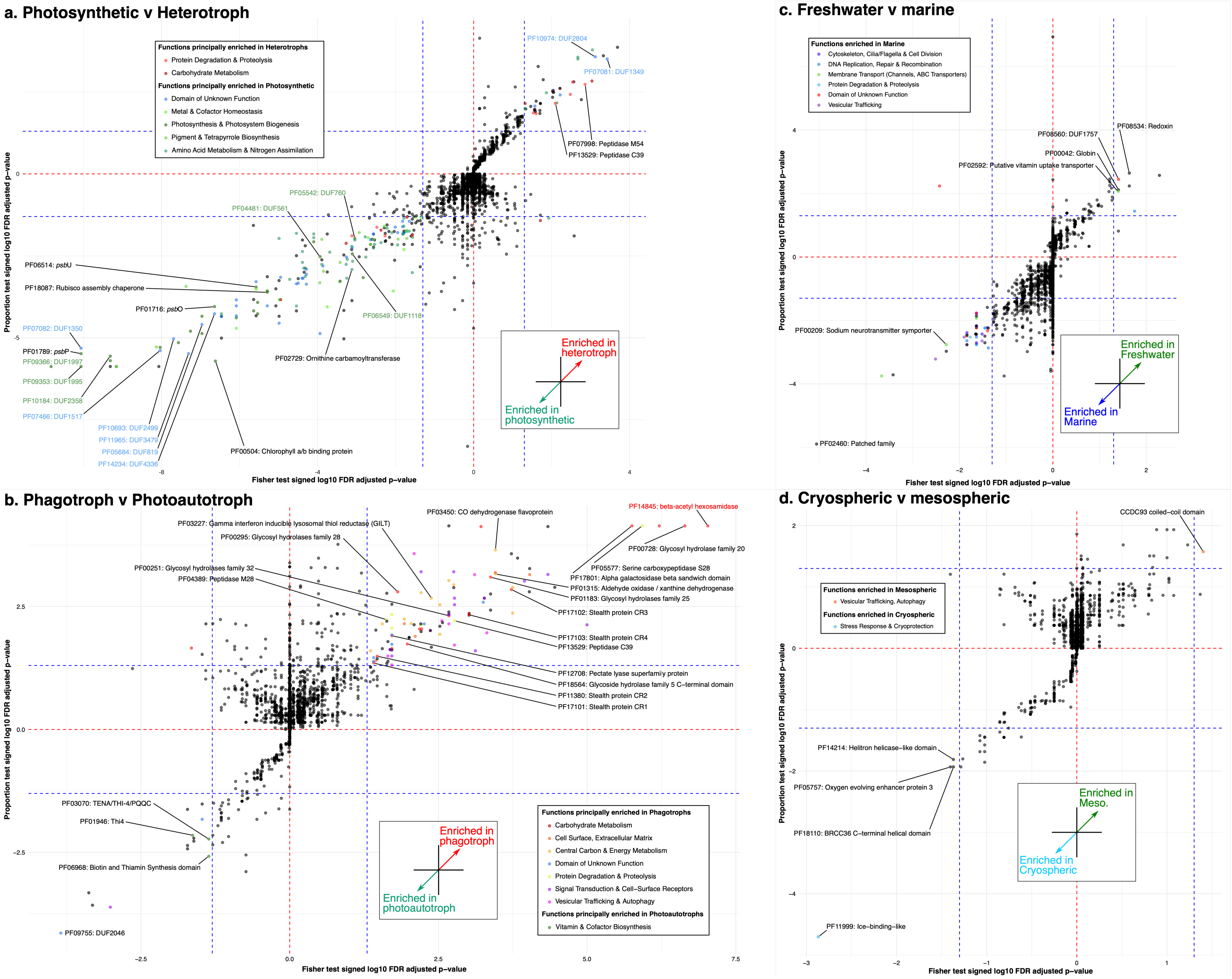
Comparative analysis of genomic composition between environmental, trophic and thermal traits of the Chrysophyceae. Each scatter plot presents the adjusted signed *p-*value of Fisher test on abscissa and adjusted signed *p-*value of Wilcoxon test of proportion on the ordinate. **a.** Comparison between photosynthetic (photoautotrophs and mixotrophs) and heterotrophic taxa (non-capable of photosynthesis). **b.** Comparison between phagotrophic (heterotrophs and mixotrophs) and photoautotrophic taxa (non-capable of phagocytosis). **c.** Comparison between freshwater (including taxa from the soil) and marine taxa. **d.** Comparison between taxa associated with the cryosphere or the mesosphere.

47 Domains of Unknown Function (DUF) were enriched in either photosynthetic (40) or heterotrophic taxa (7) and may represent novel proteins associated with trophic mode. While several photosynthesis-associated DUFs (DUF561/PF04481; DUF760/PF05542; DUF1118/PF06549; DUF1995/PF09353; DUF1997/PF09366; DUF2358/PF10184) have been inferred to have plastid-associated functions (e.g. Wakao *et al*., 2021; Yuan & Van Wijk, 2024), others have not, including some showing strong enrichment (e.g. DUF1350/PF07082). Of the heterotrophy-associated DUFs, only DUF1349 (PF07081) has external evidence of a carbohydrate-active function, adopting a concanavalin A-like lectin/glycosidase fold, and occurring fused to a GH43 glycoside hydrolase domain (also enriched in heterotrophs but above our Fisher *p*-value threshold [0.0642]) in characterized bacterial proteins (Zhang *et al*., 2019).

#### Phagotrophic versus obligately photoautotrophic taxa

119 PFAM domains were enriched either in phagotrophic (105) or photoautotrophic taxa (14), supporting net gene loss in obligate phototrophs (Figure 3b; Supplementary Table 4). The phagotroph-enriched set was dominated by functions that could be linked to phagocytic activity, including 18 domains related to carbohydrate metabolism (Supplementary Table 4), such as glycosyl hydrolases, stealth-family glycosyltransferases and the most significantly enriched domain, a beta-acetyl hexosaminidase (PF14845; P<10^-04^). Other domains enriched in phagocytotic taxa related to protein degradation and central carbon and energy metabolism (Aldehyde oxidase/xanthine-dehydrogenase-type oxidoreductase, lysosomal disulfide reductase), which may be involved in prey digestion; signal transduction and cell-surface receptors (12), possibly involved in prey detection; and vesicular trafficking and biogenesis of the digestive vacuole or late endosome (10). The photoautotrophy-enriched set was much more limited in size but strikingly included three domains relating to thiamine biosynthesis (Thi4/PF01946; biotin-thiamine synthesis-associated/PF06968; TENA-PQQC/PF03080), not detected in phagotrophic taxa (Supplementary Table 4), possibly reflecting thiamine metabolic autonomy in taxa no longer able to acquire it by phagotrophy.

#### Freshwater versus marine taxa

50 PFAMs were significantly enriched in either marine (43) or freshwater taxa (7) (Figure 3c; Supplementary Table 5). Only one has previously been linked to osmotolerance: PF00209, a sodium neurotransmitter symporter implicated in sodium-coupled ion transport under high-salinity conditions in fungi (Petrovič *et al*., 2002). The remaining marine-enriched domains spanned diverse processes, including DNA replication, repair and recombination (6), cytoskeleton, cilia and cell division (6), vesicular trafficking and organellar biogenesis (5), proteolysis (4), and 4 DUFs. The freshwater-enriched domains included two redox-related domains (redoxin/PF08534; globin/PF00042), a vitamin-uptake transporter (PF02592), and one DUF (DUF1757).

#### Mesic versus cryospheric taxa

Only 5 domains were significant (Figure 3d; Supplementary Table 6), 4 of which were enriched in cryospheric taxa: the most significant, an ice-binding-like domain (PF11999) already well-known to be associated with polar species; a helitron-helicase-like domain (PF14214); a photosystem II oxygen-evolving-enhancer (PF05757); and a BRCC36 deubiquitinase-complex protein (PF18110). A single PFAM, a CCDC93 coiled-coil domain (PF09762), was enriched in mesophilic taxa.

### Metagenome distributions of genes associated with trophism and ecology

Finally, we mapped the global marine distributions of three PFAMs: PF11999 (ice-binding domain), associated with cryosphere species, PF01716 (*psb*O) associated with photosynthesis, and PF14845 (beta-acetyl hexosaminidase) enriched in phagotrophs and absent from obligate photoautotrophs. Abundances of *Tara* Oceans unigenes (MATOUs) phylogenetically assigned to Chrysophyceae in metaT and metaG data were used to assess gene expression and abundance, respectively.

#### PF11999 - Ice binding domain

PF11999 metaT and metaG distributions were largely consistent and restricted to the Arctic (Figure 4a) with all positive stations having measured temperatures < 2.6°C. PF11999 metagene predictions were largely explained by salinity and restricted to the polar oceans, alongside the Baltic and Black Seas, which lack *Tara* data but have low salinity and may seasonally freeze (Granskog *et al*., 2006; Matov *et al*., 2022). Predicted PF11999 abundance was slightly higher for metaG than metaT data.

**Figure 4.**
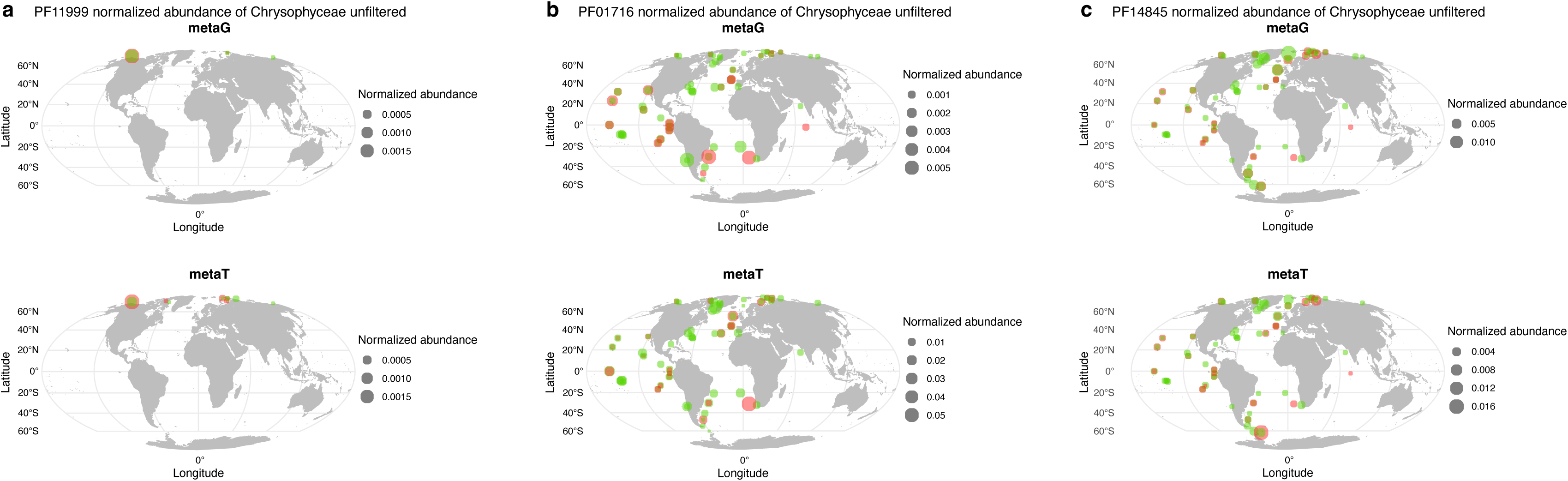
Metagenome distributions of genes associated with trophism and ecology. Mean abundances of each Chrysophyceae assigned gene at each station were normalized by the total Chrysophyceae reads at the respective station. Abundances at the surface were plotted in green and detected abundances at the DCM were plotted in red. **a.** Detected abundances in the metaG and metaT of the Chrysophyceae assigned PF11999. **b.** Detected abundances in the metaG and metaT of the Chrysophyceae assigned PF01716. **c.** Detected abundances in the metaG and metaT of the Chrysophyceae assigned PF14845.

#### PF01716 - psbO

psbO MetaT and metaG distributions diverged from one another (Figure 4b). metaT abundances were enriched in temperate and high latitude (particularly Arctic and Southern Atlantic) stations and surface depths, although much lower in the two Southern Ocean stations measured. This trend was clearest for larger size fractions (micro- and macro-), whereas pico- and nano-sizes fractions showed more ubiquitous latitudinal distributions with surface preference. In contrast, metaG distributions were more uniform across latitudes and depths. Considering environmental variables, metaT abundances were slightly negatively correlated to temperature and positively to nitrate at the surface and DCM, whereas metaG abundances were not correlated to any variable at the surface, and at the DCM were slightly positively correlated to pigments and negatively to Si and nitrate (Supplementary Figure 25).

#### PF14845 - beta-acetyl hexosaminidase

MetaT and metaG abundances were generally congruent to one another and ubiquitous (Figure 4c; Supplementary Figure 26). Both showed greater relative abundance in polar stations, predominantly in nano-size fractions, while at lower latitudes, particularly in pico-size fractions, abundances were frequently equivalent or greater at the DCM than at the surface. PF14845 abundances were positively correlated to nitrate at the surface (metaT and metaG) and DCM (metaG only), and to phosphate and oxygen at the surface, and negatively correlated to pigments at both surface (chlorophyll c3, peridinin, lutein) and DCM (zeaxanthin, hexanoylfucoxanthin, prasinoxanthin; Supplementary Figure 26).

Finally, Chrysophyceae PF14845 sequences were recovered in 4 polyphyletic gene families (Supplementary Figure 27) with different metagene distributions (Supplementary Figure 28): clade A was the least abundant and predominantly restricted to temperate/tropical latitudes; clade B was abundant and ubiquitous in metaT and metaG data; clade C was concentrated towards polar stations in metaG and lower latitudes in metaT data; and clade D was concentrated towards polar stations in metaG but homogenously distributed in metaT data.

## Discussion

We demonstrate how Chrysophyceae, historically considered a freshwater group but found ubiquitously in marine ecosystems can be used to connect algal taxonomy, ecology, and gene function. Exploiting their ecological and trophic diversity, we establish a framework to explore Chrysophycean evolution and distributions in the ocean, and the genes that underpin their ecosystem functions. These approaches are particularly instrumental to identifying novel “dark matter” genes relevant to the entire algal tree of life.

### Phylogenetic and functional diversity of Chrysophyceae

First, we present a comprehensive phylogeny of Chrysophyceae (Figure 1, covering all known orders except Apoikiida and Segregatales (Grossmann *et al*., 2016) alongside new environmental ones. This recapitulates known internal relationships beteen freshwater groups (Grossmann *et al*., 2016; Kristiansen & Škaloud, 2017; Bock *et al*., 2022; Malavasi *et al*., 2024, 2025; An *et al*., 2025; Andersen *et al*., 2025), and extends known marine Chrysophyceae diversity beyond the cultivable *Paraphysomonas* (Kristiansen & Škaloud, 2017) to multiple clades represented predominantly by MAGs, with eight transitions over the salinity barrier (Figure 1b). The polyphyly of marine Chrysophyceae is consistent with previous 18S rDNA phylogenies (Del Campo & Massana, 2011) and metagenomic evidence (Jamy *et al*., 2022).

Our dataset includes multiple strains from Arctic and Southern Oceans, and mountain glaciers, including seven independent colonizations of the cryosphere. In two cases (the sister-group to *Dinobryon*, and the order Hydrurales), freshwater and polar cryosphere strains were sister-groups, suggesting salinity transitions. For example, *Hydrurus*, associated with cold freshwater streams (Klaveness *et al*., 2011) and continental snowfields (Procházková *et al*., 2026), is sister to strains from Arctic (CCMP2296, RCC7845) and Southern Oceans (CCMP1899). Based on their growth tolerance (Perrin & Dorrell, 2024) and affinity to ice we propose the genus name *Glacidusochromonas* for the marine sister of *Hydrurus* (Figure 1), although it is itself paraphyletic, with CCMP2296 sister to all other strains. Ancestral state reconstruction suggests the order Hydrurales is likely ancestrally freshwater, colonizing polar oceans at least twice.

### Marine Chrysophyceae are ubiquitous but not uniform

Phylogenetic reconciliation of *Tara* ASVs revealed Chrysophyceae across all marine basins (Supplementary Figure 5), biased toward smaller size fractions (Supplementary Figure 6), as in previous metagenomic studies e.g. Malaspina (Obiol *et al*., 2020). Smaller Chrysophyceae might be mixotrophic or non-photosynthetic (e.g. *Paraphysomonas*), as these trophic modes are associated with smaller cell sizes (Olefeld *et al*., 2018), consistent with *psb*O being most strongly expressed in larger size fractions (Figure 4).

Different Chrysophyceae lineages showed different biogeographical patterns, including polar (Hydrurales, the sister-group to *Dinobryon*) and tropical specialists (MAG clade A, *Pedospumella*), with different determinant environmental constraints. Completing the environmental distributions of freshwater groups (Synurales, Hibberdiales) (Jeong *et al*., 2019) would require broader sampling of freshwater and soil metagenomes (Singer *et al*., 2021). In contrast, the most abundant uncultured marine groups, MAG clades A and B, which may be deep-branching free-living (Jamy *et al*., 2025) or symbiotic algae (Sørensen *et al*., 2023), will most likely be best characterized via *in situ* hybridization and microscopy of environmental samples.

### Convergent gene losses underpinning Chrysophyceae functional diversification

We infer that the Chrysophyceae ancestor was likely marine, mesic and mixotrophic (Figure 2), as are their closest relatives (Synchromophyceae, Pinguiophyceae) (Kawachi *et al*., 2002; Horn *et al*., 2007), with cryosphere colonizations, obligate photoautotrophy and obligate heterotrophy representing derived states. The large number of PFAMs associated with photosynthetic and phagocytotic life strategies indicate a driving role for gene loss in heterotrophy and photoautotrophy. Likewise, more PFAMs are associated with marine than freshwater species, which might relate to gene losses during repeated freshwater colonizations, while cold adaptation seems to be driven by a very small number of gene gains.

These approaches allowed us to identify marker genes of Chrysophyceae functional biology (Figure 3; Figure 4). This is exemplified by the ice-binding domain (PF11999; DUF3494) enriched in taxa of the cryosphere (Dorrell *et al*., 2023). PF11999 domains inhibit ice recrystallization, ensuring algal viability in freezing conditions (Vance *et al*., 2019). Previous studies have showed Chrysophyceae IBPs are upregulated at low temperatures, e.g. in the snow alga *Kremastochrysopsis* (Raymond & Remias, 2019), and their MATOUs were restricted to the Arctic and Antarctic Oceans, matching the distribution of the major cryospheric groups (i.e. Hydrurales and the sister-group to *Dinobryon*). This biogeographic signal was independent of PF11999 phylogeny, which is highly polyphyletic and driven by repeated gene transfers (Dorrell *et al*., 2023). PF11999 domains were also detected in non-cryospheric taxa from ice-inflected environments, e.g. *Pedospumella bureschii* JBL14, collected from a puddle in Lunz, Austria (Grossmann *et al*., 2016). Other occurrences outside cryosphere taxa could represent vestigial genes, or IBPs adapted to new functions such as salinity tolerance (Freyria *et al*., 2022).

Trophic transitions have much greater impacts on Chrysophyceae cells (Kristiansen & Škaloud, 2017; Olefeld *et al*., 2018) but also genome content (Dorrell *et al*., 2019). We identified two new marine losses of photosynthesis in MAG clades A and B (Figure 1), adding to 16 documented in cultured taxa (Dorrell *et al*., 2019). We further identified a new potential loss of phagotrophy in the ancestor of tChrysosacca and Hydrurales (Figure 2), alongside the documented example of the Synurales (Andersen, 2004; Kristiansen & Škaloud, 2017). Most gene losses associated with heterotrophic taxa related to plastid and photosynthetic functions (Figure 3), consistent with previous studies (Dorrell *et al*., 2019). The phototrophy-associated DUFs, some with plant homologues, are promising candidates for future functional characterization of the molecular mechanisms of photosynthesis.

Finally, gene functionsr elated to phagotrophy (Figure 3) spanned the complete sequence of prey recognition and capture through pseudopodia characteristic of the Chrysophyceae feeding behavior (Boenigk & Arndt, 2000; Jeong *et al*., 2021b), to engulfment, phagosome maturation, trafficking and digestion. These included the gamma-interferon-inducible lysosomal thiol reductase (GILT, PF03227), the only reported lysosomal enzyme able to reduce disulfide bonds in proteolysis (Arunachalam *et al*., 2000). Conversely, three thiamine (vitamin B1)-related domains were enriched in obligate photoautotrophs, notably PF03070 (*tenA*), which catalyses the salvage of a thiamine precursor in a pathway fdescribed in bacteria (Jenkins *et al*., 2007) and present in plants (Zallot *et al*., 2014). Vitamin auxotrophy has been extensively studied in algae (Croft *et al*., 2005, 2006), including in environmental contexts (Vancaester *et al*., 2020; Dorrell *et al*., 2024), but less explicitly connected to trophic mode. We suggest that phagomixotrophic Chrysophyceae obtain thiamine from digested prey, while photoautotrophs scavenge it from the environment. Strikingly, Chryophyceae seem to lack the ThiS thiamine biosynthesis pathway in their plastid proteomes (Dorrell *et al*., 2019), and this merits further investigation.

The cellular functions of these proteins await direct characterization, e.g. via reverse genetics, which to our knowledge is not yet possible in Chrysophyceae (Faktorová *et al*., 2020); comparative ‘omics of gene expression dynamics; or heterologous expression in alternative systems (e.g., yeast; bacteria; the Eustigmatophyceae alga *Nannochloropsis*) to confirm their cellular localization and kinetics.

### Ecological roles of the marine Chrysophyceae

Finally, we connected Chrysophyceae taxonomy, trophic diversity, and ecology by mapping meta-genome distributions of *psb*O (photosynthesis) and PF14845 (phagocytosis). Some of the meta-gene distributions may relate to the biogeography of Chrysophyceaè with different trophic modes, similarly as observed in other algae (Rihm *et al*., 2025). For example, non-photosynthetic Paraphysomonadida lacking *psbO* are more abundant in tropical open oceans and subpolar waters, as observed for other small phagomixotrophic plankton (Sommeria-Klein *et al*., 2021), whereas photoautotrophic Hydrurales (McKenrie *et al*., 1995) lacking PF14845 are confined to high latitudes. Nevertheless, the most abundant, predominantly mixotrophic MAG clades A and B likely drive the total meta-gene abundances we observed.

Both gene families were detected globally (metaG) but had contrasting distributions in expression (metaT; Figure 4). *psbO* sshowed greater relative occurrence in low-latitude stations, whereas PF14845 showed greater relative abundance in high-latitude stations. In contrast, we observed a greater investment in *psb*O expression in metaT stations from high latitudes, large size fractions, and surface depths; whereas PF14845 was more occurrent in metaT data from DCM and low latitude samples. Finally, *psb*O metaT abundances were negatively correlated to temperature at the surface, PF14845 negatively to pigments in the DCM, and both positively to nitrate at both depths.

Together, the metaT and metaG distributions place temperature as the most structuring ecological variable of Chrysophyceae trophic status. The ubiquity of *psb*O and PF14845 suggests that Chrysophyceae are capable of both photosynthesis and phagocytosis globally, but the poise between the two shifts between environments. In nutrient-rich, illuminated surface waters at high latitudes, photosynthesis predominates, including in photoautotrophic groups such as Hydrurales. In temperate waters photomixotrophy likely predominates, with both photosynthesis and heterotrophy stimulated by nutrients. In oligotrophic, tropical, and deep-water environments, obligate heterotrophy, e.g. bacterivory, likely predominates. Trophic poise may also fluctuate seasonally, e.g. with heterotrophy predominating in light-limited polar winters or during blooms of competitors or prey, which may explain the low *psb*O metaT abundances in Southern Ocean *Tara* stations.

This tradeoff has implications in the context of global change. Warming oceans would favour low-latitude, warm-adapted, largely mixotrophic and heterotrophic Chrysophyceae in lieu of high-latitude, cold-adapted photoautotrophs. Experimental studies of mixotrophic protists, including Chrysophyceae, consistently show that warming shifts mixotrophic metabolism towards heterotrophy (Wilken *et al*., 2020; Lepori-Bui *et al*., 2022; Barbaglia *et al*., 2024; Moeller *et al*., 2024; Honig *et al*., 2025), a shift maintained, or even amplified by acidification (Slomka *et al*., 2026). At the community level, global surveys of *Tara* Oceans plankton diversity and interaction networks similarly project that warming will disproportionately restructure polar plankton communities as temperate and tropical species migrate to higher latitudes (Ibarbalz *et al*., 2019; Chaffron *et al*., 2021; Frémont *et al*., 2022). Our results suggest that ocean warming will reshape marine Chrysophyceae along two connected axes: a poleward loss of cold-adapted photoautotrophic groups such as Hydrurales, and an expansion of warm-adapted mixotrophic MAG lineages, who will increasingly rely on heterotrophy.

## Acknowledgements

Work in the Dorrell Group is supported by an ERC Starting Grant, ChloroMosaic (101037960), awarded to RGD 2023-2028. The authors thank Connie Lovejoy (Université Laval) for the provision of *Glacidusochromonas* sp. CCMP2296. The authors thank Daniel Vaulot (Station Biologique de Roscoff) for the provision of the marine Chrysophyceae strains RCC6673, RCC6679. The authors thank the Undergraduate and Masters interns that supported the making of this study, Sarah Tonnelier (Université Paris Sciences Lettres, Paris, France) and Tyler Desharnais (California State University, San Marcos, USA).

## Competing interests

The authors declare no competing interests.

## Author contributions

L.G. and R.G.D. designed the project. L.G., I.P. and R.G.D. assembled and maintained the strains collection. L.G. and R.G.D. conducted the experiments (RNA extractions). L.G., E.Z., X.Z. and R.G.D. designed the bioinformatics analysis analyzed and interpreted the data. L.G. and R.G.D. wrote the draft and all authors read and agreed on the manuscript.

## Data availability

All R codes used to generate maps, PCA and predicted abundances were made available on GitHub (https://github.com/Dorrell-group/Chrysophyceae-EcoGenomicFramework/tree/main). Supporting material and data were deposited on FigShare (https://doi.org/10.6084/m9.figshare.34814514).

## Notes

### Competing Interest Statement

The authors have declared no competing interest.

## References

Ali RH, Bogusz M, Whelan S. 2019. Identifying Clusters of High Confidence Homologies in Multiple Sequence Alignments (K Tamura, Ed.). Molecular Biology and Evolution 36: 2340–2351.

An Y, Hao J, Nan F, Lv J, Liu Q, Liu X, Xie S, Feng J. 2025. New records of the colonial chrysophyte Urostipulosphaera articulata (Chrysophyceae, Ochromonadales) in China. Biodiversity Data Journal 13: e150900.

Andersen RA. 2004. Biology and systematics of heterokont and haptophyte algae. American Journal of Botany 91: 1508–1522.

Andersen RA, Graf L, Malakhov Y, Kim H, Yoon HS. 2025. *Stylococcus aureus* and *S. brevis sp. nov.* (Chrysophyceae) and their phylogenetic relationship to *Lagynion*. Phycologia 64: 282–296.

Arunachalam B, Phan UT, Geuze HJ, Cresswell P. 2000. Enzymatic reduction of disulfide bonds in lysosomes: Characterization of a Gamma-interferon-inducible lysosomal thiol reductase (GILT). Proceedings of the National Academy of Sciences 97: 745–750.

Barbaglia GS, Paight C, Honig M, Johnson MD, Marczak R, Lepori-Bui M, Moeller HV. 2024. Environment-dependent metabolic investments in the mixotrophic chrysophyte *Ochromonas*. Journal of Phycology 60: 170–184.

Beisser D, Graupner N, Bock C, Wodniok S, Grossmann L, Vos M, Sures B, Rahmann S, Boenigk J. 2017. Comprehensive transcriptome analysis provides new insights into nutritional strategies and phylogenetic relationships of chrysophytes. PeerJ 5: e2832.

Benjamini Y, Hochberg Y. 1995. Controlling the false discovery rate: A practical and powerful approach to multiple testing. Journal of the Royal Statistical Society Series B: Statistical Methodology 57: 289–300.

Berges JA, Franklin DJ, Harrison PJ. 2001. Evolution of an artificial seawater medium: Improvements in enriched seawater, artificial water over the last two decades. Journal of Phycology 37: 1138–1145.

Bjorbækmo MFM, Evenstad A, Røsæg LL, Krabberød AK, Logares R. 2020. The planktonic protist interactome: where do we stand after a century of research? The ISME Journal 14: 544–559.

Bock C, Olefeld JL, Vogt JC, Albach DC, Boenigk J. 2022. Phylogenetic and functional diversity of Chrysophyceae in inland waters. Organisms Diversity & Evolution 22: 327–341.

Boden L, Klagus C, Boenigk J. 2024. Insights from single-strain and mixed culture experiments on the effects of heatwaves on freshwater flagellates. PeerJ 12: e17912.

Boenigk J, Arndt H. 2000. Particle Handling during Interception Feeding by Four Species of Heterotrophic Nanoflagellates. Journal of Eukaryotic Microbiology 47: 350–358.

Boyer TP, García HE, Locarnini RA, Zweng MM, Mishonov AV, Reagan JR, Weathers KA, Baranova OK, Paver CR, Seidov D, et al. 2018. World Ocean Atlas.

Brown JW, Sorhannus U. 2010.A molecular genetic timescale for the diversification of autotrophic Stramenopiles (Ochrophyta): substantive underestimation of putative fossil ages (MTP Gilbert, Ed.). PLoS ONE 5: e12759.

Büdel B, Friedl T, Beyschlag W (Eds). 2024. Biology of Algae, Lichens and Bryophytes. Berlin, Heidelberg: Springer Berlin Heidelberg.

Burns JA, Pittis AA, Kim E. 2018. Gene-based predictive models of trophic modes suggest Asgard archaea are not phagocytotic. Nature Ecology & Evolution 2: 697–704.

Bushmanova E, Antipov D, Lapidus A, Prjibelski AD. 2019. rnaSPAdes: a *de novo* transcriptome assembler and its application to RNA-Seq data. GigaScience 8: giz100.

Busi SB, Bourquin M, Fodelianakis S, Michoud G, Kohler TJ, Peter H, Pramateftaki P, Styllas M, Tolosano M, De Staercke V, et al. 2022. Genomic and metabolic adaptations of biofilms to ecological windows of opportunity in glacier-fed streams. Nature Communications 13: 2168.

Capella-Gutiérrez S, Silla-Martínez JM, Gabaldón T. 2009. trimAl: a tool for automated alignment trimming in large-scale phylogenetic analyses. Bioinformatics 25: 1972–1973.

Caron DA, Alexander H, Allen AE, Archibald JM, Armbrust EV, Bachy C, Bell CJ, Bharti A, Dyhrman ST, Guida SM, et al. 2017. Probing the evolution, ecology and physiology of marine protists using transcriptomics. Nature Reviews Microbiology 15: 6–20.

Caron DA, Countway PD, Jones AC, Kim DY, Schnetzer A. 2012. Marine Protistan Diversity. Annual Review of Marine Science 4: 467–493.

Caron DA, Worden AZ, Countway PD, Demir E, Heidelberg KB. 2009. Protists are microbes too: a perspective. The ISME Journal 3: 4–12.

Chaffron S, Delage E, Budinich M, Vintache D, Henry N, Nef C, Ardyna M, Zayed AA, Junger PC, Galand PE, et al. 2021. Environmental vulnerability of the global ocean epipelagic plankton community interactome. Science Advances 7: eabg1921.

Chen S. 2025.fastp 1.0: An ultra-fast all-round tool for FASTQ data quality control and preprocessing. iMeta 4: e70078.

Choi S-W, Graf L, Choi JW, Jo J, Boo GH, Kawai H, Choi CG, Xiao S, Knoll AH, Andersen RA, et al. 2024. Ordovician origin and subsequent diversification of the brown algae. Current Biology 34: 740–754.e4.

Croft MT, Lawrence AD, Raux-Deery E, Warren MJ, Smith AG. 2005. Algae acquire vitamin B12 through a symbiotic relationship with bacteria. Nature 438: 90–93.

Croft MT, Warren MJ, Smith AG. 2006. Algae Need Their Vitamins. Eukaryotic Cell 5: 1175–1183.

De Vargas C, Audic S, Henry N, Decelle J, Mahé F, Logares R, Lara E, Berney C, Le Bescot N, Probert I, et al. 2015. Eukaryotic plankton diversity in the sunlit ocean. Science 348: 1261605.

Del Campo J, Massana R. 2011. Emerging diversity within Chrysophytes, Choanoflagellates and Bicosoecids based on molecular surveys. Protist 162: 435–448.

Delmont TO, Gaia M, Hinsinger DD, Frémont P, Vanni C, Fernandez-Guerra A, Eren AM, Kourlaiev A, d’Agata L, Clayssen Q, et al. 2022. Functional repertoire convergence of distantly related eukaryotic plankton lineages abundant in the sunlit ocean. Cell Genomics 2: 100123.

Dorrell RG, Azuma T, Nomura M, Audren De Kerdrel G, Paoli L, Yang S, Bowler C, Ishii K, Miyashita H, Gile GH, et al. 2019. Principles of plastid reductive evolution illuminated by nonphotosynthetic chrysophytes. Proceedings of the National Academy of Sciences 116: 6914–6923.

Dorrell RG, Gile G, McCallum G, Méheust R, Bapteste EP, Klinger CM, Brillet-Guéguen L, Freeman KD, Richter DJ, Bowler C. 2017. Chimeric origins of ochrophytes and haptophytes revealed through an ancient plastid proteome. eLife 6: e23717.

Dorrell RG, Kuo A, Füssy Z, Richardson EH, Salamov A, Zarevski N, Freyria NJ, Ibarbalz FM, Jenkins J, Pierella Karlusich JJ, et al. 2023. Convergent evolution and horizontal gene transfer in Arctic Ocean microalgae. Life Science Alliance 6: e202201833.

Dorrell RG, Nef C, Altan-Ochir S, Bowler C, Smith AG. 2024. Presence of vitamin B_12_ metabolism in the last common ancestor of land plants. Philosophical Transactions of the Royal Society B: Biological Sciences 379: 20230354.

Eddy SR. 2011.Accelerated Profile HMM Searches (WR Pearson, Ed.). PLoS Computational Biology 7: e1002195.

Faktorová D, Nisbet RER, Fernández Robledo JA, Casacuberta E, Sudek L, Allen AE, Ares M, Aresté C, Balestreri C, Barbrook AC, et al. 2020. Genetic tool development in marine protists: emerging model organisms for experimental cell biology. Nature Methods 17: 481–494.

Falkowski P. 2012. Ocean science: The power of plankton. Nature 483: S17–S20.

Falkowski PG, Fenchel T, Delong EF. 2008. The microbial engines that drive Earth’s biogeochemical cycles. Science 320: 1034–1039.

Frémont P, Gehlen M, Vrac M, Leconte J, Delmont TO, Wincker P, Iudicone D, Jaillon O. 2022. Restructuring of plankton genomic biogeography in the surface ocean under climate change. Nature Climate Change 12: 393–401.

Frey W. 2020. Syllabus of Plant Families - A. Engler’s Syllabus der Pflanzenfamilien Part 2/1: Glaucocystophyta, Cryptophyta, Dinophyta/Dinozoa, Haptophyta, Heterokontophyta/Ochrophyta, Chlorarachniophyta/Cercozoa, Euglenophyta/Euglenozoa, Chlorophyta, Streptophyta p.p. Stuttgart: Schweizerbart.

Freyria NJ, Kuo A, Chovatia M, Johnson J, Lipzen A, Barry KW, Grigoriev IV, Lovejoy C. 2022. Salinity tolerance mechanisms of an Arctic Pelagophyte using comparative transcriptomic and gene expression analysis. Communications Biology 5: 500.

Friedlingstein P, O’Sullivan M, Jones MW, Andrew RM, Bakker DCE, Hauck J, Landschützer P, Le Quéré C, Luijkx IT, Peters GP, et al. 2023. Global Carbon Budget 2023. Earth System Science Data 15: 5301–5369.

Gasol JM, Kirchman DL (Eds). 2018. Microbial ecology of the oceans. Hoboken, NJ: Wiley Blackwell.

Gérikas Ribeiro C, Dos Santos AL, Gourvil P, Le Gall F, Marie D, Tragin M, Probert I, Vaulot D. 2020. Culturable diversity of Arctic phytoplankton during pack ice melting (JW Deming and C Michel, Eds). Elementa: Science of the Anthropocene 8: 6.

Granskog M, Kaartokallio H, Kuosa H, Thomas DN, Vainio J. 2006. Sea ice in the Baltic Sea – A review. Estuarine, Coastal and Shelf Science 70: 145–160.

Grigoriev IV, Hayes RD, Calhoun S, Kamel B, Wang A, Ahrendt S, Dusheyko S, Nikitin R, Mondo SJ, Salamov A, et al. 2021. PhycoCosm, a comparative algal genomics resource. Nucleic Acids Research 49: D1004–D1011.

Grossmann L, Bock C, Schweikert M, Boenigk J. 2016. Small but Manifold – Hidden Diversity in “ *Spumella* -like Flagellates”. Journal of Eukaryotic Microbiology 63: 419–439.

Guiry MD, Guiry GM. 2025. AlgaeBase. *World-wide electronic publication, University of Galway*.

Guschina IA, Harwood JL. 2006. Lipids and lipid metabolism in eukaryotic algae. Progress in Lipid Research 45: 160–186.

Hoang DT, Chernomor O, Von Haeseler A, Minh BQ, Vinh LS. 2018. UFBoot2: Improving the Ultrafast Bootstrap Approximation. Molecular Biology and Evolution 35: 518–522.

Honig MA, Barbaglia GS, Doyle MD, Moeller HV. 2025. Effects of mixotroph evolution on trophic transfer (BE Beisner, Ed.). Journal of Plankton Research 47: fbae053.

Horn S, Ehlers K, Fritzsch G, Gil-Rodríguez MC, Wilhelm C, Schnetter R. 2007. Synchroma grande spec. nov. (Synchromophyceae class. nov., Heterokontophyta): An Amoeboid Marine Alga with Unique Plastid Complexes. Protist 158: 277–293.

Ibarbalz FM, Henry N, Brandão MC, Martini S, Busseni G, Byrne H, Coelho LP, Endo H, Gasol JM, Gregory AC, et al. 2019. Global Trends in Marine Plankton Diversity across Kingdoms of Life. Cell 179: 1084–1097.e21.

Jamy M, Biwer C, Vaulot D, Obiol A, Jing H, Peura S, Massana R, Burki F. 2022. Global patterns and rates of habitat transitions across the eukaryotic tree of life. Nature Ecology & Evolution 6: 1458–1470.

Jamy M, Huber T, Antoine T, Ruscheweyh H-J, Paoli L, Pelletier E, O. Delmont T, Burki F. 2025. Identification of a deep-branching lineage of algae using environmental plastid genomes. Nature Communications 17: 662.

Jenkins AH, Schyns G, Potot S, Sun G, Begley TP. 2007. A new thiamin salvage pathway. Nature Chemical Biology 3: 492–497.

Jeong HJ, Kang HC, Lim AS, Jang SH, Lee K, Lee SY, Ok JH, You JH, Kim JH, Lee KH, et al. 2021a. Feeding diverse prey as an excellent strategy of mixotrophic dinoflagellates for global dominance. Science Advances 7: eabe4214.

Jeong M, Kim JI, Jo BY, Kim HS, Siver PA, Shin W. 2019. Surviving the marine environment: two new species of *Mallomonas* (Synurophyceae). Phycologia 58: 276–286.

Jeong M, Kim JI, Nam SW, Shin W. 2021b. Molecular phylogeny and taxonomy of the genus *Spumella* (Chrysophyceae) based on morphological and molecular evidence. Frontiers in Plant Science 12: 758067.

Jones P, Binns D, Chang H-Y, Fraser M, Li W, McAnulla C, McWilliam H, Maslen J, Mitchell A, Nuka G, et al. 2014. InterProScan 5: genome-scale protein function classification. Bioinformatics 30: 1236–1240.

Josse J, Husson F. 2016. missMDA: A package for handling missing values in multivariate data analysis. Journal of Statistical Software 70.

Kamiya M. 2020. Syllabus of Plant Families - A. Engler’s Syllabus der Pflanzenfamilien Part 2/2. Stuttgart: Schweizerbart.

Katoh K, Rozewicki J, Yamada KD. 2019. MAFFT online service: multiple sequence alignment, interactive sequence choice and visualization. Briefings in Bioinformatics 20: 1160–1166.

Kawachi M, Inouye I, Honda D, O’Kelly CJ, Bailey JC, Bidigare RR, Andersen RA. 2002. The Pinguiophyceae classis nova, a new class of photosynthetic stramenopiles whose members produce large amounts of omega-3 fatty acids. Phycological Research 50: 31–47.

Keeling PJ, Burki F, Wilcox HM, Allam B, Allen EE, Amaral-Zettler LA, Armbrust EV, Archibald JM, Bharti AK, Bell CJ, et al. 2014.The Marine Microbial Eukaryote Transcriptome Sequencing Project (MMETSP): Illuminating the Functional Diversity of Eukaryotic Life in the Oceans through Transcriptome Sequencing (RG Roberts, Ed.). PLoS Biology 12: e1001889.

Keller MD, Selvin RC, Claus W, Guillard RRL. 1987. Media for the culture of oceanic ultraphytoplankton. Journal of Phycology 23: 633–638.

Kozlov AM, Darriba D, Flouri T, Morel B, Stamatakis A. 2019. RAxML-NG: a fast, scalable and user-friendly tool for maximum likelihood phylogenetic inference (J Wren, Ed.). Bioinformatics 35: 4453–4455.

Kristiansen J, Škaloud P. 2017. Chrysophyta. In: Archibald JM, Simpson AGB, Slamovits CH, eds. Handbook of the Protists. Cham: Springer International Publishing, 331–366.

Lang I, Hodac L, Friedl T, Feussner I. 2011. Fatty acid profiles and their distribution patterns in microalgae: a comprehensive analysis of more than 2000 strains from the SAG culture collection. BMC Plant Biology 11: 124.

Larsson A. 2014. AliView: a fast and lightweight alignment viewer and editor for large datasets. Bioinformatics 30: 3276–3278.

Lepori-Bui M, Paight C, Eberhard E, Mertz CM, Moeller HV. 2022. Evidence for evolutionary adaptation of mixotrophic nanoflagellates to warmer temperatures. Global Change Biology 28: 7094–7107.

Letunic I, Bork P. 2024. Interactive Tree of Life (iTOL) v6: recent updates to the phylogenetic tree display and annotation tool. Nucleic Acids Research 52: W78–W82.

Levering J, Broddrick J, Dupont CL, Peers G, Beeri K, Mayers J, Gallina AA, Allen AE, Palsson BO, Zengler K. 2016.Genome-Scale Model Reveals Metabolic Basis of Biomass Partitioning in a Model Diatom (A Ianora, Ed.). PLOS ONE 11: e0155038.

Li-Beisson Y, Thelen JJ, Fedosejevs E, Harwood JL. 2019. The lipid biochemistry of eukaryotic algae. Progress in Lipid Research 74: 31–68.

Malavasi V, Pusztai M, Jadrná I, Škvorová Z, Škaloud P. 2024. Morphological diversity and phylogeny of the palmelloid chrysophyte genera *Chrysotilos* and *Globulochrysis*, gen. nov. European Journal of Phycology 59: 279–289.

Malavasi V, Pusztai M, Jankowska K, Zakryś B, Škaloud P. 2025. Revisiting Chrysococcus (Chrysophyceae): new phylogenetic evidence and evolutionary implications. Fottea 25: 121–127.

Manni M, Berkeley MR, Seppey M, Simão FA, Zdobnov EM. 2021. BUSCO Update: Novel and Streamlined Workflows along with Broader and Deeper Phylogenetic Coverage for Scoring of Eukaryotic, Prokaryotic, and Viral Genomes (J Kelley, Ed.). Molecular Biology and Evolution 38: 4647–4654.

Matov M, Peneva E, Galabov V. 2022. Black Sea Freezing and Relation to the Winter Conditions in 2006–2021. Atmosphere 13: 974.

Maurer-Alcalá XX, Kim E. 2024. TIdeS: A Comprehensive Framework for Accurate Open Reading Frame Identification and Classification in Eukaryotic Transcriptomes (J Archibald, Ed.). Genome Biology and Evolution 16: evae252.

McKenrie CH, Deibel D, Paranjape MA, Thompson RJ. 1995. The marine mixotroph *Dinobryon balticum* (Chrysophyceae): phagotrophy and survival in a cold ocean. Journal of Phycology 31: 19–24.

Mistry J, Chuguransky S, Williams L, Qureshi M, Salazar GA, Sonnhammer ELL, Tosatto SCE, Paladin L, Raj S, Richardson LJ, et al. 2021. Pfam: The protein families database in 2021. Nucleic Acids Research 49: D412–D419.

Moeller HV, Archibald KM, Leles SG, Pfab F. 2024. Predicting optimal mixotrophic metabolic strategies in the global ocean. Science Advances 10: eadr0664.

Nonoyama T, Kazamia E, Nawaly H, Gao X, Tsuji Y, Matsuda Y, Bowler C, Tanaka T, G. Dorrell R. 2019. Metabolic Innovations Underpinning the Origin and Diversification of the Diatom Chloroplast. Biomolecules 9: 322.

Novák Vanclová AM, Nef C, Füssy Z, Vancl A, Liu F, Bowler C, Dorrell RG. 2024. New plastids, old proteins: repeated endosymbiotic acquisitions in kareniacean dinoflagellates. EMBO Reports 25: 16.

Obiol A, Giner CR, Sánchez P, Duarte CM, Acinas SG, Massana R. 2020. A metagenomic assessment of microbial eukaryotic diversity in the global ocean. Molecular Ecology Resources 20: 718–731.

Olefeld JL, Majda S, Albach DC, Marks S, Boenigk J. 2018. Genome size of chrysophytes varies with cell size and nutritional mode. Organisms Diversity & Evolution 18: 163–173.

Paradis E, Schliep K. 2019. ape 5.0: an environment for modern phylogenetics and evolutionary analyses in R (R Schwartz, Ed.). Bioinformatics 35: 526–528.

Penot-Raquin M, Novák Vanclová AMG, Powell V, Corbeau Y, Younes C, Eugène M, Bouceba T, Pionneau C, De Almeida Bastos V, Garcia M, et al. 2026. Characterisation of the new microalgal protein xATPA related to the F-type ATP synthase α subunit, from the ecosystem to the molecule.

Perrin AJ, Dorrell RG. 2024. Protists and protistology in the Anthropocene: challenges for a climate and ecological crisis. BMC Biology 22: 279.

Pesant S, Not F, Picheral M, Kandels-Lewis S, Le Bescot N, Gorsky G, Iudicone D, Karsenti E, Speich S, Troublé R, et al. 2015. Open science resources for the discovery and analysis of Tara Oceans data. Scientific Data 2: 150023.

Petrovič U, Gunde-Cimerman N, Plemenitaš A. 2002. Cellular responses to environmental salinity in the halophilic black yeast *Hortaea werneckii*. Molecular Microbiology 45: 665–672.

Pierella Karlusich JJ, Cosnier K, Zinger L, Henry N, Nef C, Bernard G, Scalco E, Dvorak E, Tara Oceans Coordinators, Acinas SG, et al. 2025. Patterns and drivers of diatom diversity and abundance in the global ocean. Nature Communications 16: 3452.

Pierella Karlusich JJ, Pelletier E, Zinger L, Lombard F, Zingone A, Colin S, Gasol JM, Dorrell RG, Henry N, Scalco E, et al. 2023. A robust approach to estimate relative phytoplankton cell abundances from metagenomes. Molecular Ecology Resources 23: 16–40.

Pous S, Lazure P, Carton X. 2015. A model of the general circulation in the Persian Gulf and in the Strait of Hormuz: Intraseasonal to interannual variability. Continental Shelf Research 94: 55–70.

Price MN, Dehal PS, Arkin AP. 2009. FastTree: Computing Large Minimum Evolution Trees with Profiles instead of a Distance Matrix. Molecular Biology and Evolution 26: 1641–1650.

Procházková L, Andersen RA, Leya T, Řezanka T, Lukeš M, Nedbalová L, Remias D. 2026. Novel *Hydrurus* species (Chrysophyceae) and their adaptations to high-altitude European and Arctic snowfields. Journal of Phycology 62: 818–845.

R Core Team. 2025. R: A language and environment for statistical computing. R Foundation for Statistical Computing, *Vienna, Austria*.

Raymond JA, Remias D. 2019. Ice-Binding Proteins in a Chrysophycean Snow Alga: Acquisition of an Essential Gene by Horizontal Gene Transfer. Frontiers in Microbiology 10: 2697.

Rihm G, Benedetti F, Bittner L. 2025. Do trophic strategies shape biogeography and environmental niches? Marine dinoflagellates as a case study. ISME Communications 5: ycaf153.

Sigee DC. 2005. Freshwater microbiology: biodiversity and dynamic interactions of microorganisms in the aquatic environment. Hoboken, NJ: J. Wiley.

Singer D, Seppey CVW, Lentendu G, Dunthorn M, Bass D, Belbahri L, Blandenier Q, Debroas D, De Groot GA, De Vargas C, et al. 2021. Protist taxonomic and functional diversity in soil, freshwater and marine ecosystems. Environment International 146: 106262.

Slomka S, Verspagen JMH, Rodriguez-Perez RB, Huisman J, Wilken S. 2026. Changes in photosynthesis and grazing facilitate growth of a mixotrophic protist under ocean acidification and warming. New Phytologist 251: 2574–2587.

Sommeria-Klein G, Watteaux R, Ibarbalz FM, Pierella Karlusich JJ, Iudicone D, Bowler C, Morlon H. 2021. Global drivers of eukaryotic plankton biogeography in the sunlit ocean. Science 374: 594–599.

Sørensen MES, Zlatogursky VV, Onuţ-Brännström I, Walraven A, Foster RA, Burki F. 2023. A novel kleptoplastidic symbiosis revealed in the marine centrohelid Meringosphaera with evidence of genetic integration. Current Biology 33: 3571–3584.e6.

Steinegger M, Söding J. 2017. MMseqs2 enables sensitive protein sequence searching for the analysis of massive data sets. Nature Biotechnology 35: 1026–1028.

Stiller JW, Schreiber J, Yue J, Guo H, Ding Q, Huang J. 2014. The evolution of photosynthesis in chromist algae through serial endosymbioses. Nature Communications 5: 5764.

Tice AK, Žihala D, Pánek T, Jones RE, Salomaki ED, Nenarokov S, Burki F, Eliáš M, Eme L, Roger AJ, et al. 2021. PhyloFisher: A phylogenomic package for resolving eukaryotic relationships (A Hejnol, Ed.). PLOS Biology 19: e3001365.

Vancaester E, Depuydt T, Osuna-Cruz CM, Vandepoele K. 2020. Comprehensive and Functional Analysis of Horizontal Gene Transfer Events in Diatoms (FU Battistuzzi, Ed.). Molecular Biology and Evolution 37: 3243–3257.

Vance TDR, Bayer-Giraldi M, Davies PL, Mangiagalli M. 2019. Ice-binding proteins and the ‘domain of unknown function’ 3494 family. The FEBS Journal 286: 855–873.

Vaulot D, Sim CWH, Ong D, Teo B, Biwer C, Jamy M, Lopes Dos Santos A. 2022. METAPR^2^ : A database of eukaryotic 18S RRNA metabarcodes with an emphasis on protists. Molecular Ecology Resources 22: 3188–3201.

Vernette C, Henry N, Lecubin J, De Vargas C, Hingamp P, Lescot M. 2021. The Ocean barcode atlas: A web service to explore the biodiversity and biogeography of marine organisms. Molecular Ecology Resources 21: 1347–1358.

Wakao S, Shih PM, Guan K, Schackwitz W, Ye J, Patel D, Shih RM, Dent RM, Chovatia M, Sharma A, et al. 2021.Discovery of photosynthesis genes through whole-genome sequencing of acetate-requiring mutants of Chlamydomonas reinhardtii (M Schroda, Ed.). PLOS Genetics 17: e1009725.

Whelan S, Irisarri I, Burki F. 2018. PREQUAL: detecting non-homologous characters in sets of unaligned homologous sequences (J Hancock, Ed.). Bioinformatics 34: 3929–3930.

Wideman JG, Monier A, Rodríguez-Martínez R, Leonard G, Cook E, Poirier C, Maguire F, Milner DS, Irwin NAT, Moore K, et al. 2019. Unexpected mitochondrial genome diversity revealed by targeted single-cell genomics of heterotrophic flagellated protists. Nature Microbiology 5: 154–165.

Wilken S, Choi CJ, Worden AZ. 2020. Contrasting Mixotrophic Lifestyles Reveal Different Ecological Niches in Two Closely Related Marine Protists. Journal of Phycology 56: 52–67.

Wong T, Ly-Trong N, Ren H, Baños H, Roger A, Susko E, Bielow C, De Maio N, Goldman N, Hahn M, et al. 2025. IQ-TREE 3: Phylogenomic Inference Software using Complex Evolutionary Models.

Worden AZ, Follows MJ, Giovannoni SJ, Wilken S, Zimmerman AE, Keeling PJ. 2015. Rethinking the marine carbon cycle: Factoring in the multifarious lifestyles of microbes. Science 347: 1257594.

Worden AZ, Nolan JK, Palenik B. 2004. Assessing the dynamics and ecology of marine picophytoplankton: The importance of the eukaryotic component. Limnology and Oceanography 49: 168–179.

Yuan B, Van Wijk KJ. 2024. The chloroplast protease system degrades stromal DUF760-1 and DUF760-2 domain-containing proteins at different rates. Plant Physiology 196: 1788–1801.

Zallot R, Yazdani M, Goyer A, Ziemak MJ, Guan J-C, McCarty DR, De Crécy-Lagard V, Gerdes S, Garrett TJ, Benach J, et al. 2014. Salvage of the thiamin pyrimidine moiety by plant TenA proteins lacking an active-site cysteine. Biochemical Journal 463: 145–155.

Zhang R, Li N, Liu Y, Han X, Tu T, Shen J, Xu S, Wu Q, Zhou J, Huang Z. 2019. Biochemical and structural properties of a low-temperature-active glycoside hydrolase family 43 β-xylosidase: Activity and instability at high neutral salt concentrations. Food Chemistry 301: 125266.

