## Supplementary Figures 1-28 for "Chaos and repetition in algal evolution: A Chrysophyceae eco-genomic framework": SupplementaryFigure_1.pdf

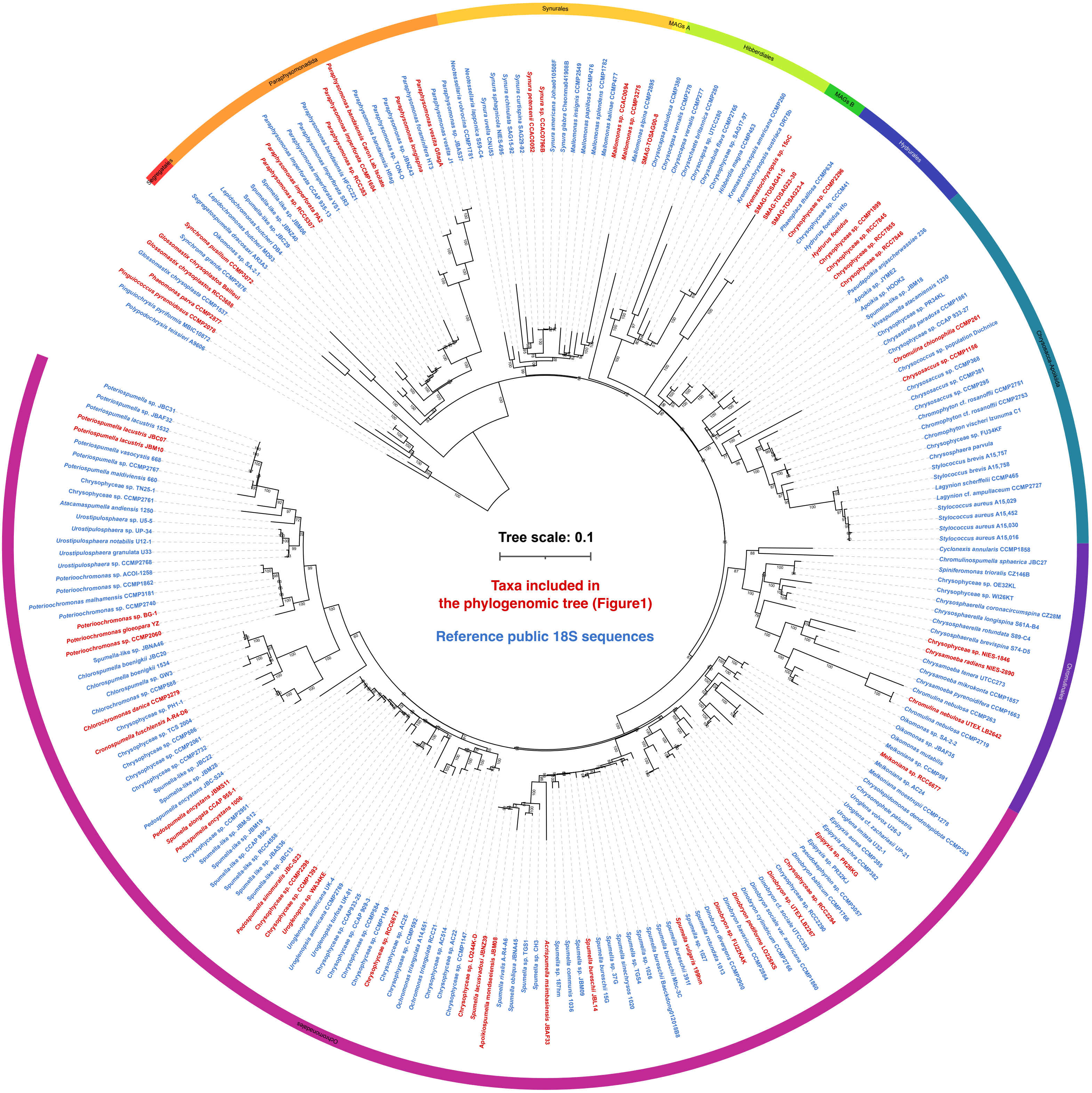


**Supplementary Figure 1.** Maximum likelihood phylogenetic tree reconstructed with18S sequences of the cultivable diversity of the Chrysophyceae. The tree was inferred with IQ-Tree 3.1.1 using the GTR+F+R6 evolutionary model. Ultrafast Bootstrap Approximation are shown near the nodes. Taxa in red are included in the phylogenomic tree and the tree search was constrained by the topology of the phylogenomic tree from Figure 1.
