## Supplementary Figures 1-28 for "Chaos and repetition in algal evolution: A Chrysophyceae eco-genomic framework": SupplementaryFigure_2.pdf

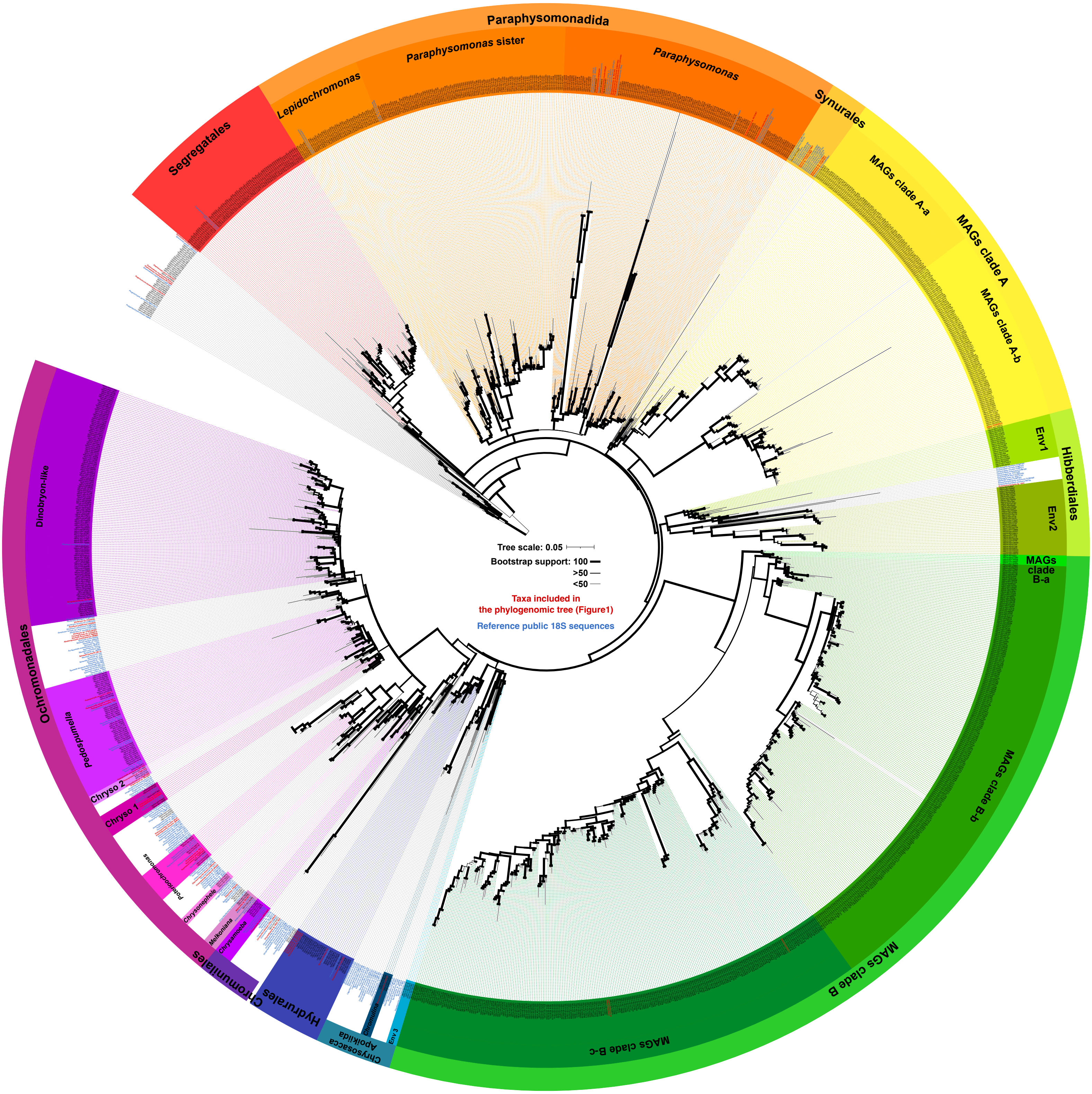


**Supplementary Figure 2.** Maximum likelihood phylogenetic tree reconstructed with18S sequences of the cultivable diversity of the Chrysophyceae (in blue and red) and the *Tara* v4 ASVs (in black). The tree was inferred with IQ-Tree 3.1.1 using the GTR+F+I+R8 evolutionary model. Ultrafast Bootstrap Approximation are shown as branch thickness. The tree search was constrained by the topology of the tree in Supplementary Figure 1.
