## Supplementary Figures 1-28 for "Chaos and repetition in algal evolution: A Chrysophyceae eco-genomic framework": SupplementaryFigure_3.pdf

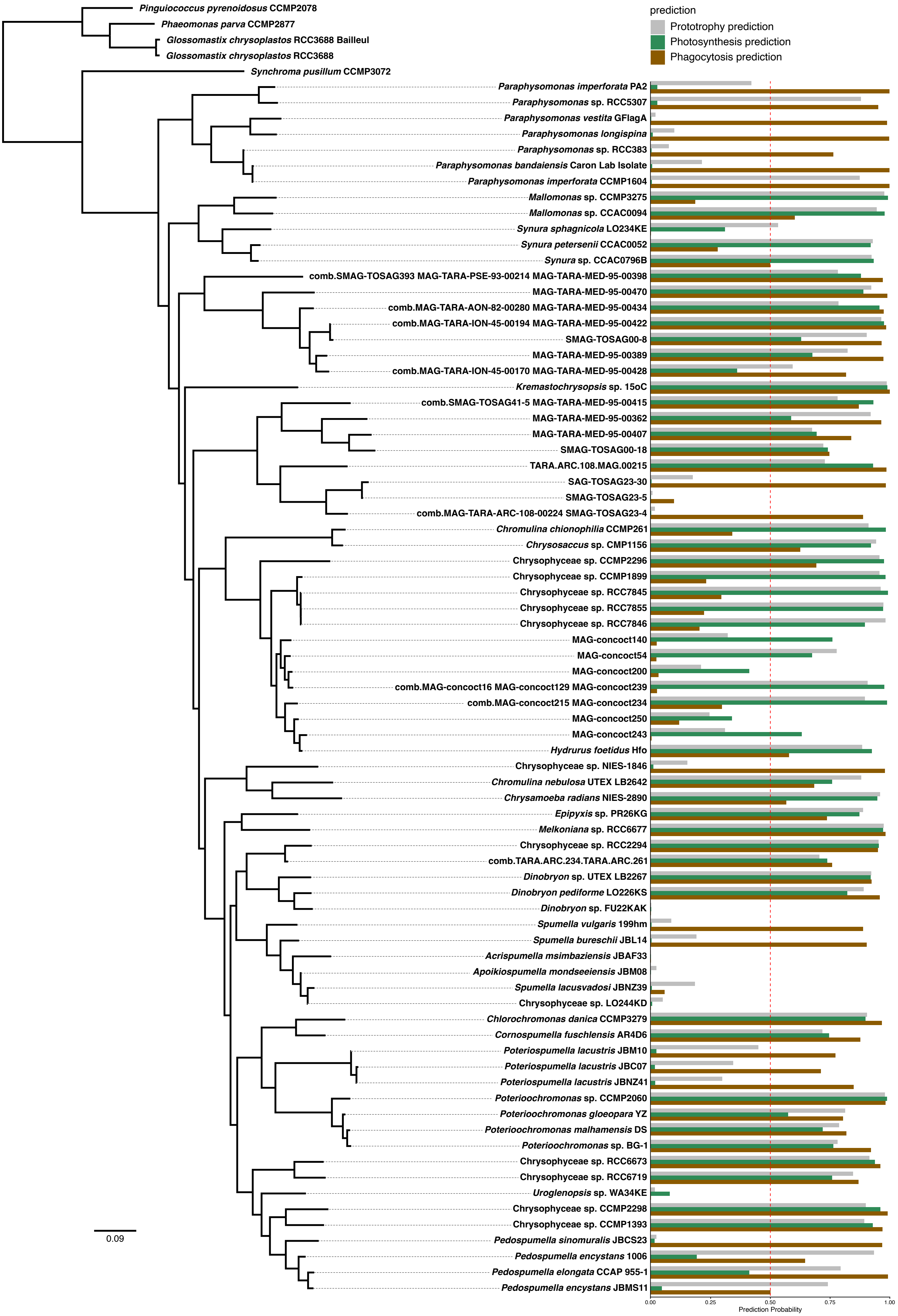


**Supplementary Figure 3.** Mapping of the probability of the different trophic modes (prototrophy, photosynthesis and phagotrophy) predicted by the model from Burns et al. 2026. The prediction were mapped on the topology of the phylogenomic tree with taxa combined from Figure 1.
