## Supplementary Figures 1-28 for "Chaos and repetition in algal evolution: A Chrysophyceae eco-genomic framework": SupplementaryFigure_4.pdf

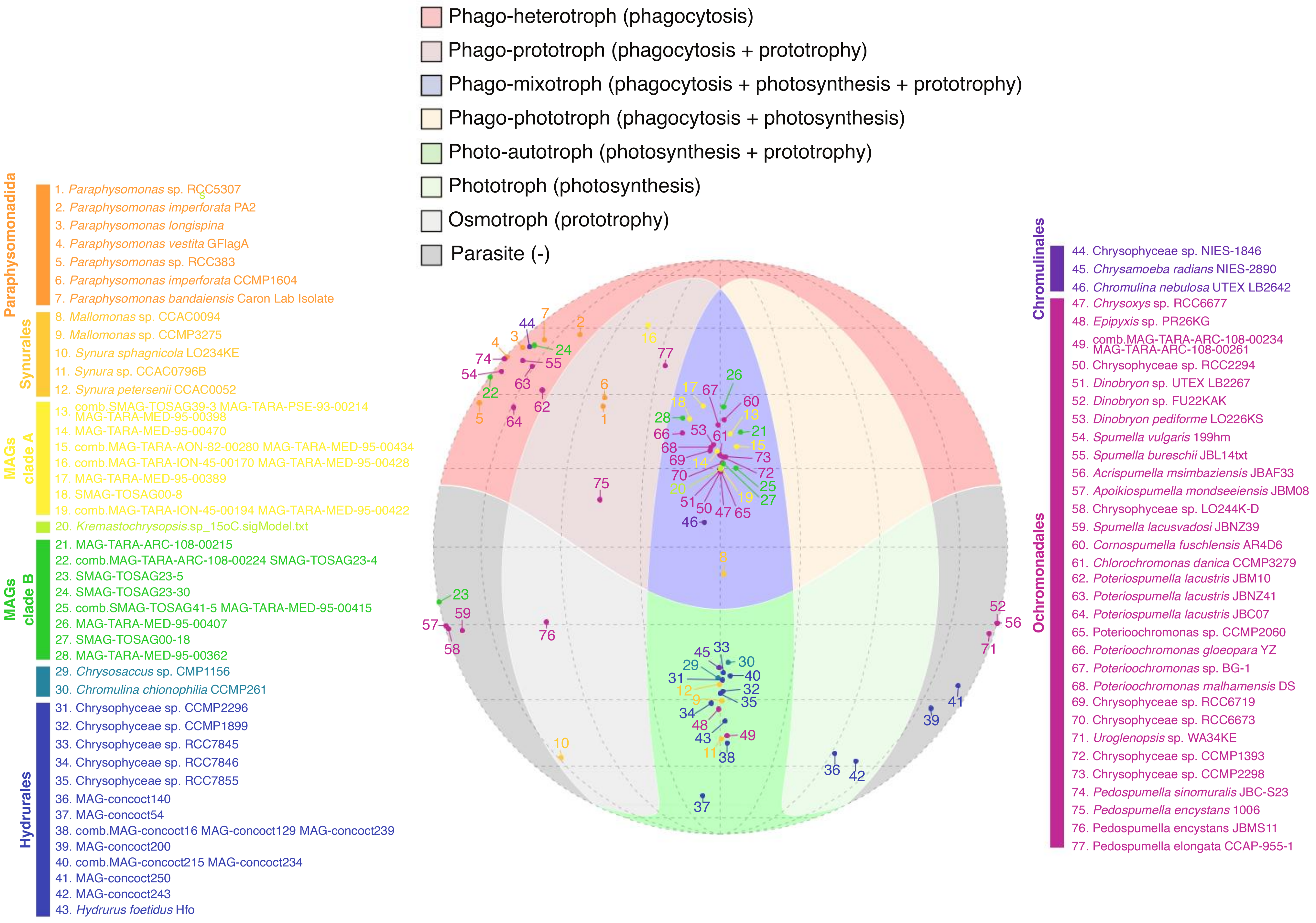


**Supplementary Figure 4.** Mollweide projection showing the combination of trophic modes predictions (i.e prototrophy, photosynthesis and phagotrophy). Each color represent a specific trophic mode indicated in the legend above the plot. Each dot indicates the taxa tested.
