## Supplementary Figures 1-28 for "Chaos and repetition in algal evolution: A Chrysophyceae eco-genomic framework": SupplementaryFigure_5.pdf

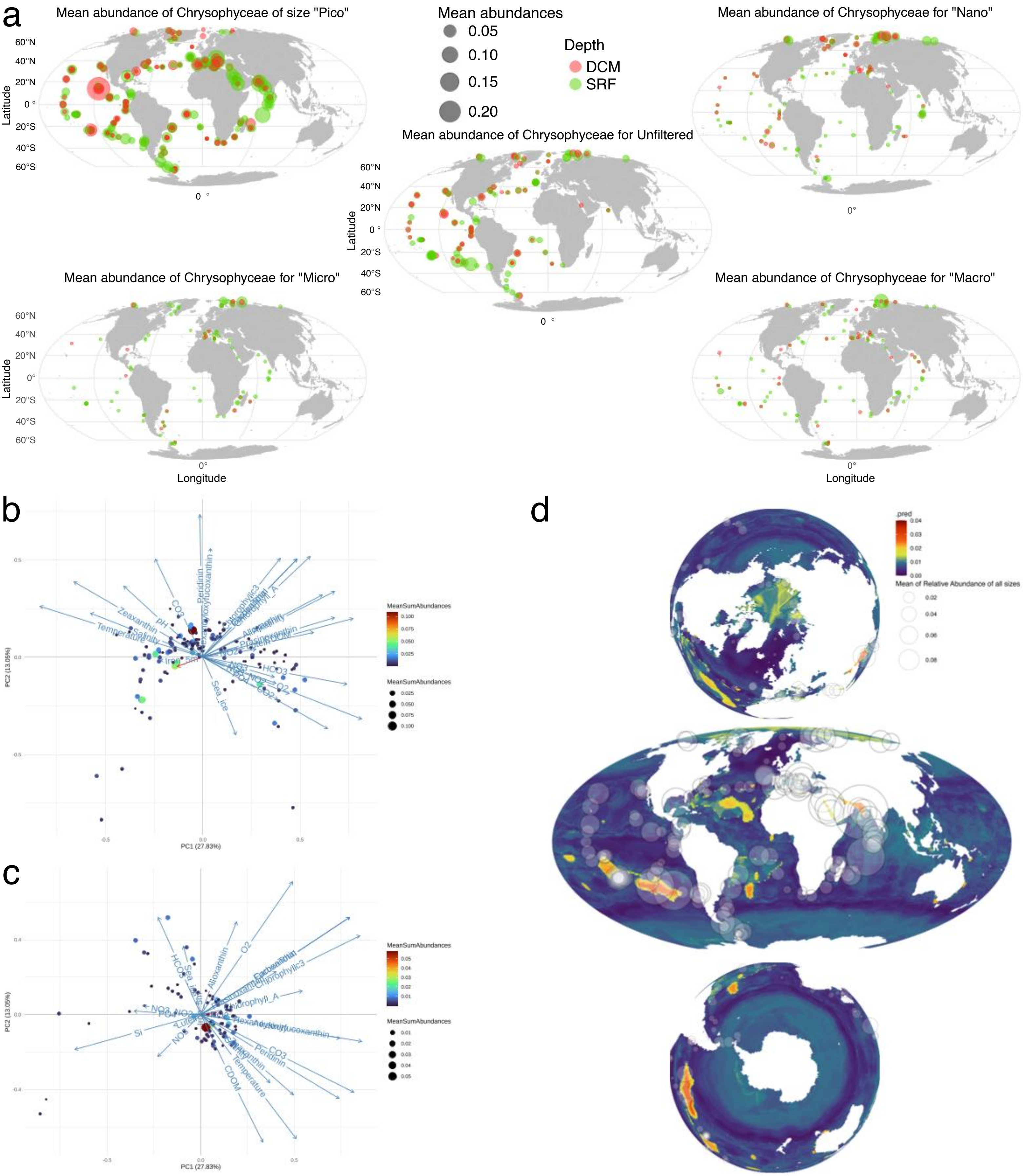


**Supplementary Figure 5.** Metabarcode informed biogeography of the Chrysophyceae. **a.** Observed mean abundances at different filter sizes. The "Pico" fraction ranged from 0.22-5$\mu$m; "Nano" from 0.8-20$\mu$m; "Micro" from 20-180$\mu$m; "Macro" from 180-2000$\mu$m; "Unfiltered" without filtration. **b.** PCA of the physicho-chemical variables in all *Tara* stations at the surface. Each dot represents a station and their respective mean abundances were represented by size and coloring. **c.** PCA of the physicho-chemical variables in all *Tara* stations at the DCM. Each dot represents a station and their respective mean abundances were represented by size and coloring. **d.** Predicted and observed abundances at the surface. All maps present the same predicted abundances but in different projections. Circles represent the observed abundances.
