## Supplementary Figures 1-28 for "Chaos and repetition in algal evolution: A Chrysophyceae eco-genomic framework": SupplementaryFigure_8.pdf

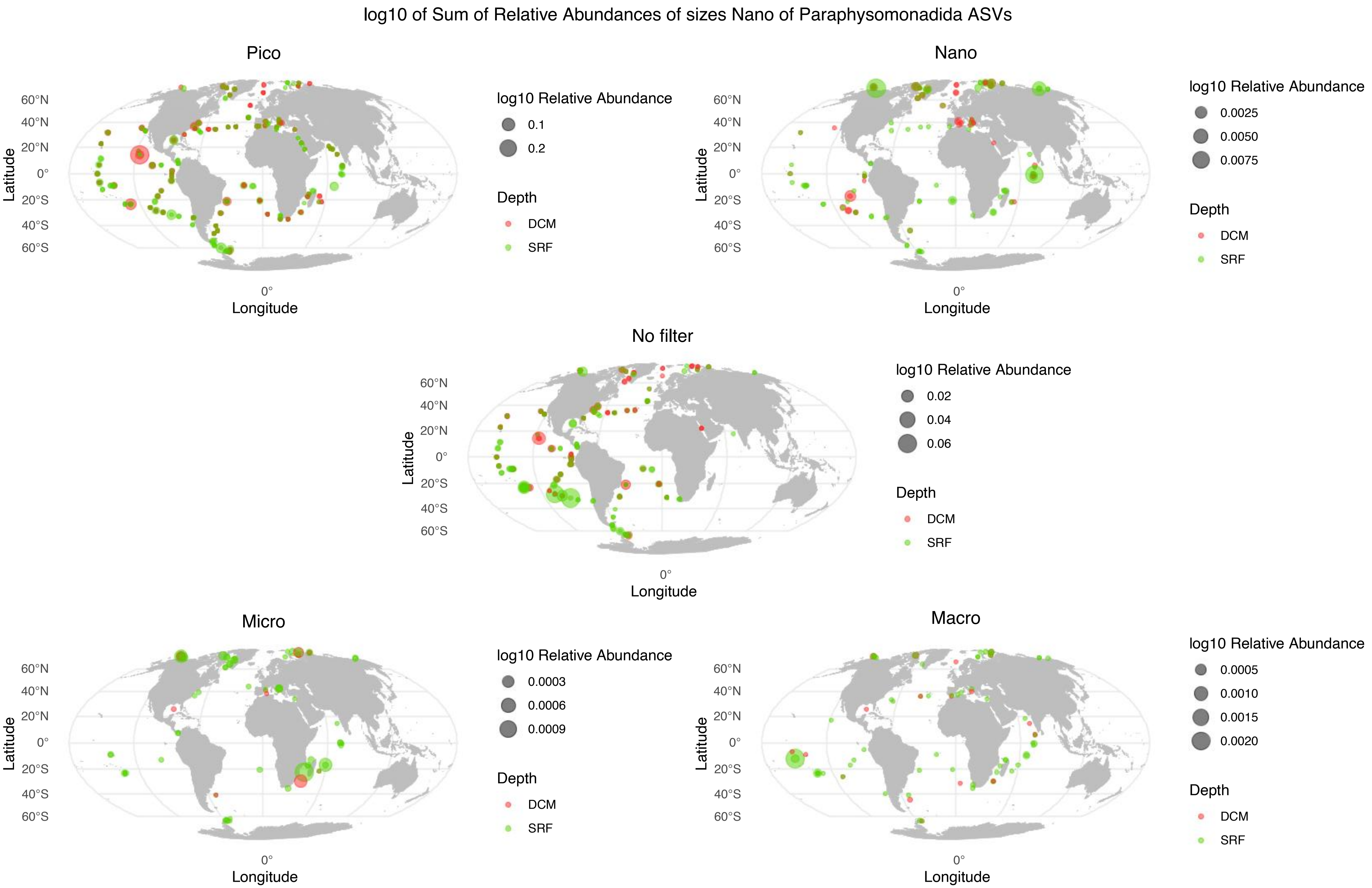


**Supplementary Figure 8.** Observed abundances of metabarcode assigned to the Paraphysomonadida lineage, at the surface (green) and the DCM (red). The abundances were separated by size fractions, "Pico" fraction ranged from 0.22-5$\mu m$; "Nano" from 0.8-20$\mu m$; "Micro" from 20-180$\mu m$; "Macro" from 180-2000$\mu m$.
