## Supplementary Figures 1-28 for "Chaos and repetition in algal evolution: A Chrysophyceae eco-genomic framework": SupplementaryFigure_21.pdf

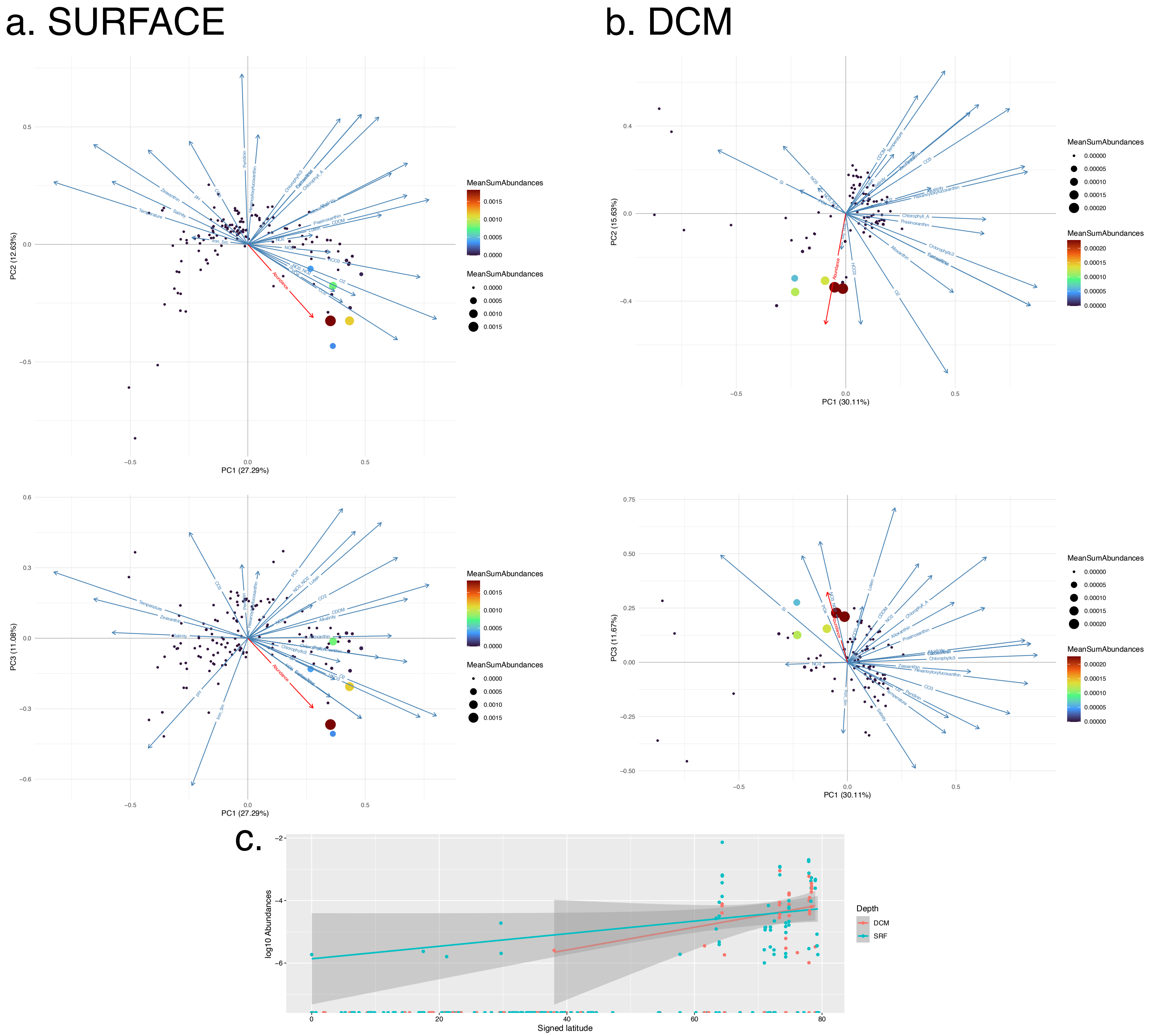


**Supplementary Figure 21.** Physico-chemical variables correlations to observed abundances of metabarcode assigned to the Hydrurales lineage. **a.** PCA of the physicho-chemical variables in all *Tara* stations at the surface. Each dot represents a station and their respective mean abundances were represented by size and coloring. **b.** PCA of the physicho-chemical variables in all *Tara* stations at the DCM. Each dot represents a station and their respective mean abundances were represented by size and coloring. **c.** Plot of the log10 abundances as a function of signed latitude ranging from 0 at the Equator to 80 near the Poles, at the Surface (cyan) and at the DCM (red).
