## Supplementary Figures 1-28 for "Chaos and repetition in algal evolution: A Chrysophyceae eco-genomic framework": SupplementaryFigure_23.pdf

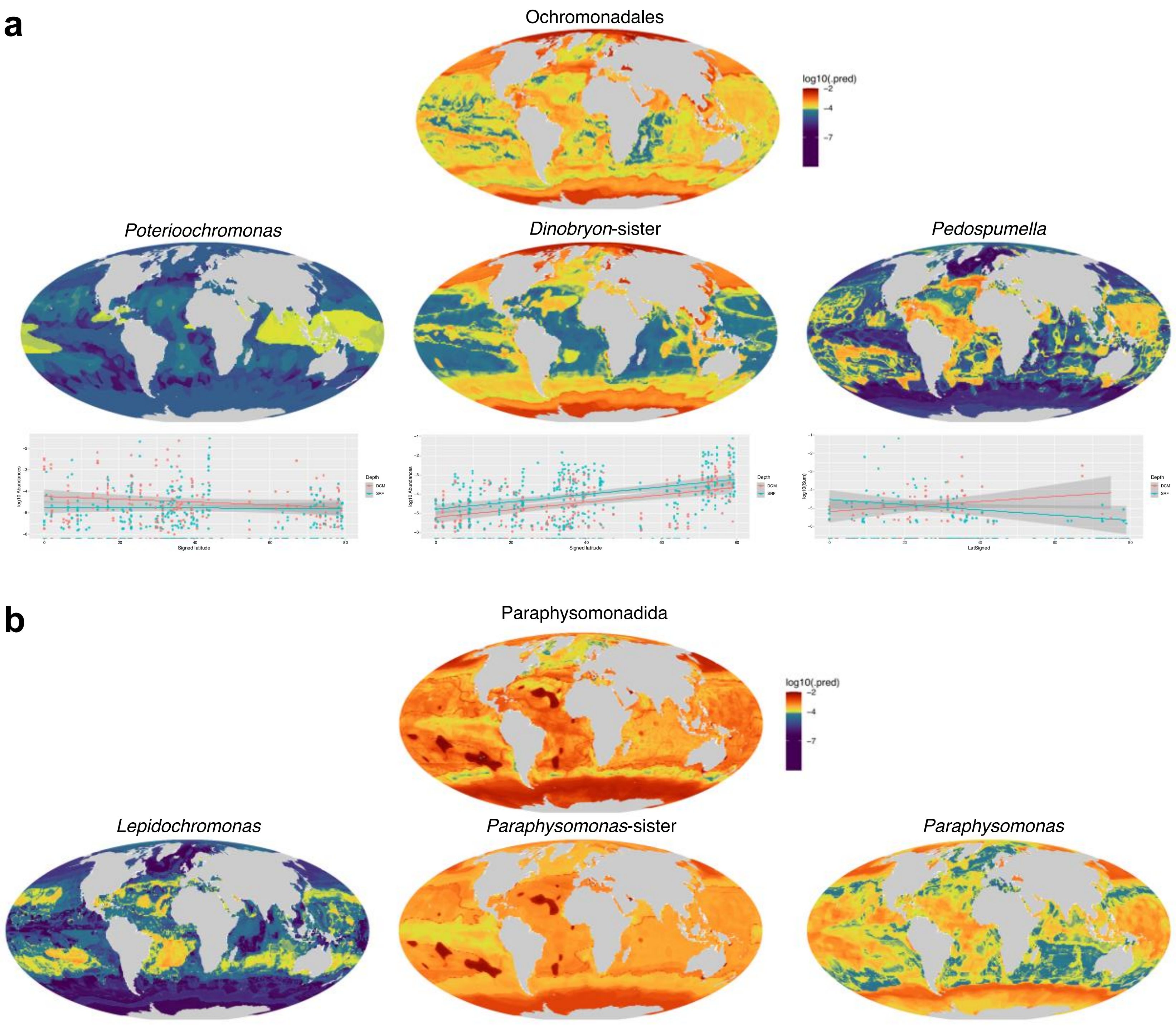


**Supplementary Figure 23.** Sublineages predicted distributions **a.** Sublineages of the Ochromonadales; *Poterioochromonas*; *Dinobryon*-sister; *Pedospumella*. Under each predictions map, we plotted of the log10 abundances as a function of signed latitude ranging from 0 at the Equator to 80 near the Poles, at the Surface (cyan) and at the DCM (red). **b.** Sublineages of the Paraphysomonadida; *Lepidochromonas*; *Paraphysomonas*-sister; *Paraphysomonas*.
