## Supplementary Figures 1-28 for "Chaos and repetition in algal evolution: A Chrysophyceae eco-genomic framework": SupplementaryFigure_25.pdf

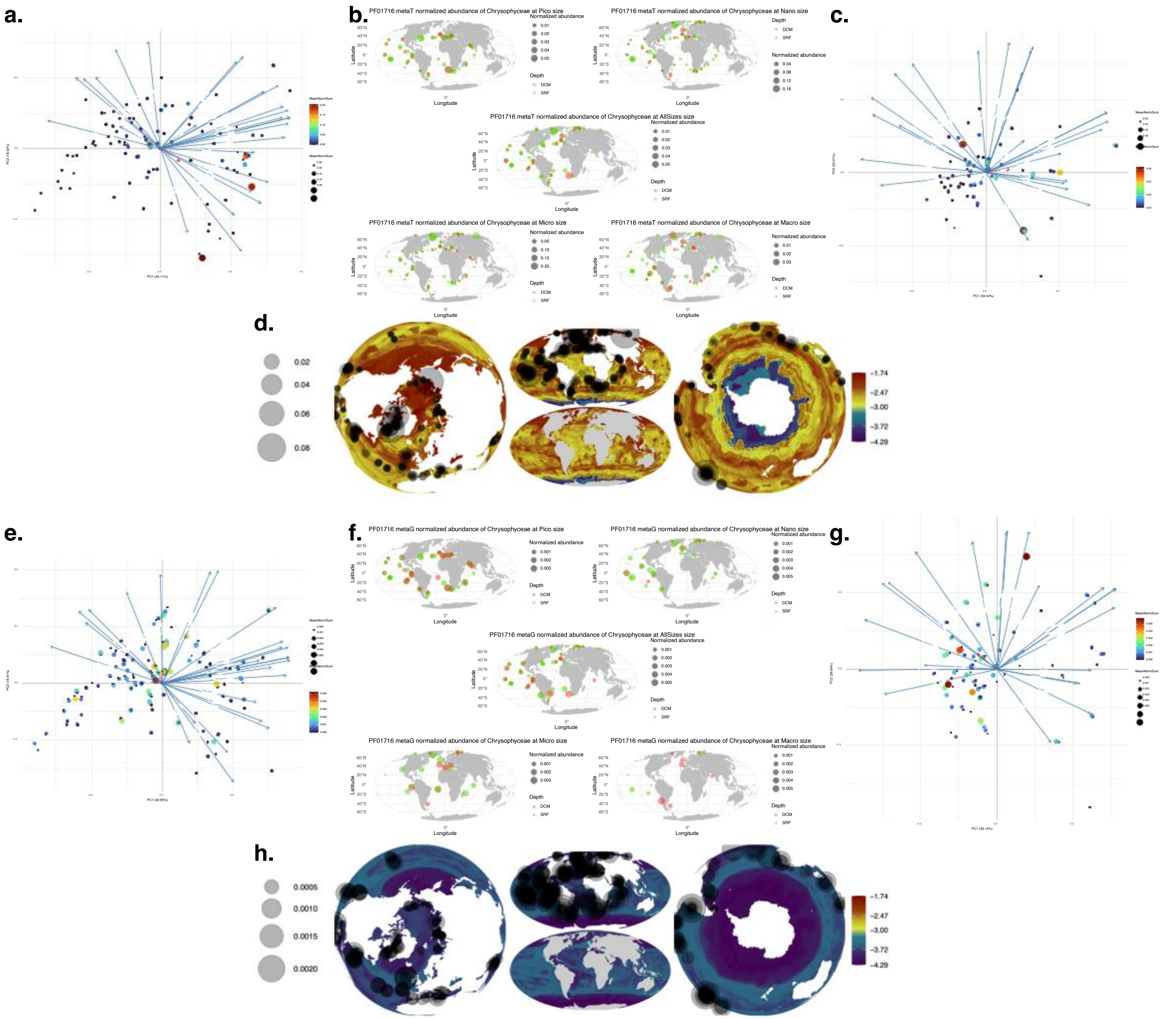


**Supplementary Figure 25.** Metagenomic distribution of observed and predicted abundances and physico-chemical variables correlation to observed abundances of the ice-binding protein PF01716 taxonomically assigned to the Chrysophyceae. **a.** PCA of the physicho-chemical variables in all *Tara* stations to metaT observed abundances at the surface. Each dot represents a station and their respective mean abundances were represented by size and coloring. **b.** Observed metaT abundances at the surface (green) and the DCM (red). The abundances were separated by size fractions, "Pico" fraction ranged from 0.22-5$\mu$m; "Nano" from 0.8-20$\mu$m; "Micro" from 20-180$\mu$m; "Macro" from 180-2000$\mu$m. **c.** PCA of the physicho-chemical variables in all *Tara* stations to metaT observed abundances at the DCM. Each dot represents a station and their respective mean abundances were represented by size and coloring. **d.** Distribution of log10 metaT predicted abundances. **e.** PCA of the physicho-chemical variables in all *Tara* stations to metaG observed abundances at the surface. Each dot represents a station and their respective mean abundances were represented by size and coloring. **f.** Observed metaG abundances at the surface (green) and the DCM (red). The abundances were separated by size fractions, "Pico" fraction ranged from 0.22-5$\mu$m; "Nano" from 0.8-20$\mu$m; "Micro" from 20-180$\mu$m; "Macro" from 180-2000$\mu$m. **g.** PCA of the physicho-chemical variables in all *Tara* stations to metaG observed abundances at the DCM. Each dot represents a station and their respective mean abundances were represented by size and coloring. **h.** Distribution of log10 metaG predicted abundances.
