## Supplementary Figures 1-28 for "Chaos and repetition in algal evolution: A Chrysophyceae eco-genomic framework": SupplementaryFigure_28.pdf

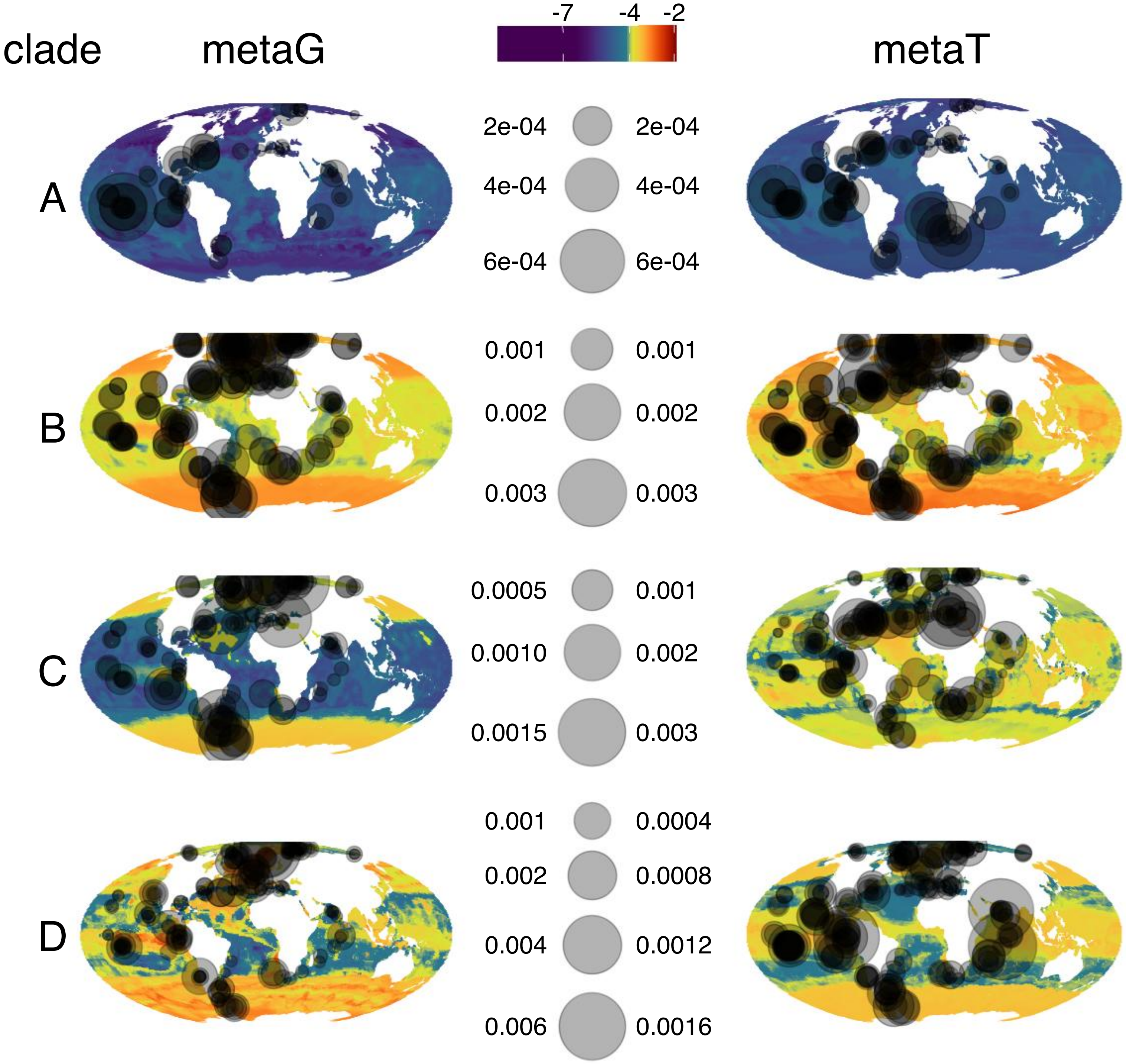


**Supplementary Figure 28. Predicted abundances of the metaT and metaG in the four clades of the Chrysophyceae PF14845.** MATOUs of the Chrysophyceae were phylogenetically assigned to the different clades (Supplementary Figure 27) and their relative abundances at the surface in the metaT and metaG were used to predict distributions using a random tree forest approach and satellite data. **a.** Clade A. **b.** Clade B. **c.** Clade C. **d.** Clade D.
