## Supplementary figures and images for "Chaos and repetition in algal evolution: A Chrysophyceae eco-genomic framework"

### SupplementaryFigure_27.pdf

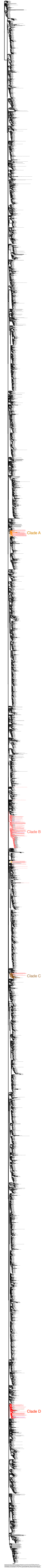
